## Supplementary material for "Cis-regulatory evolution spotlights species differences in the adaptive potential of gene expression plasticity": Fig S1-S13 and Table S1-S9

### Supplementary information

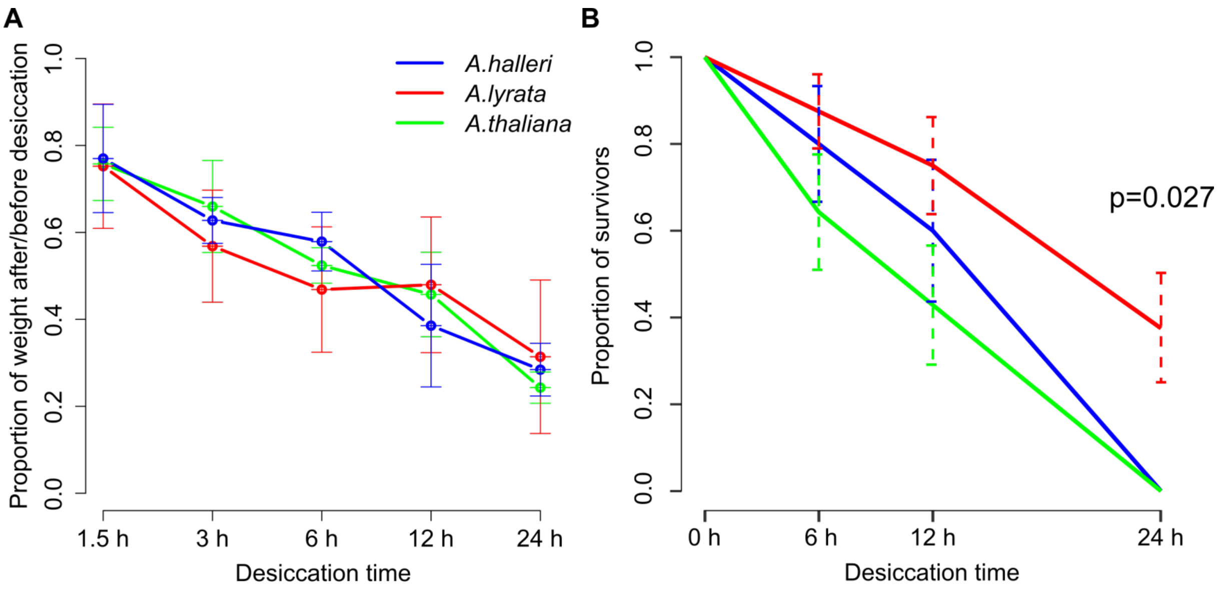


Fig. S1: (A) **The rate of water loss after 1.5 to 24h of desiccation is similar between the species.** The rate of water loss was inferred by the ratio of plant weight after / before the desiccation stress. Samples were weighted before the initiation of stress and immediately prior to RNA sampling. Species did not differ significantly in their rate of water loss (error bars indicate standard deviation, Interaction term species:duration of dehydration , F=0.562, *p*=0.803, glm model). Data shown in Supplementary Data 1.

(B) ***A. lyrata* is the most resistant to acute dehydration.** We performed the same dehydration experiment as for the transcriptome analysis, but instead of sampling material, we repotted the plants after 0h, 6h, 12h and 24h of dehydration and measured survival after 18 days. The survival rate after stress among species was significantly different and after 24h of dehydration, only *A. lyrata* survived (interaction term species:duration of dehydration in glm model, $F_{6,111}$= 8.71, *p*= 8.841e-08). This experiment included a representative set of 10, 17 and 14 genotypes of *A. halleri, A. lyrata* and *A. thaliana*, respectively (Table S8). Data shown in Supplementary Data 2.

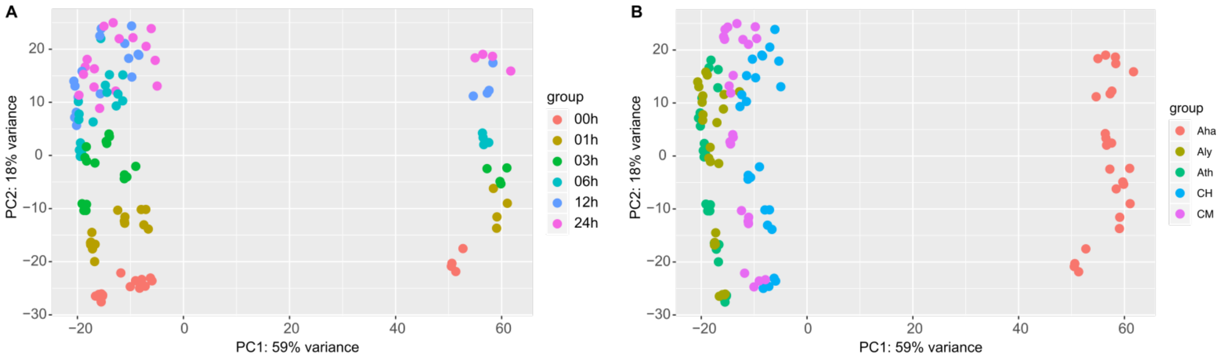


Figure S2. **Differences between species and time points have a clear genetic basis.** Principal component analysis (PCA) of the 120 transcriptome samples collected over a desiccation time course for three species and two of their hybrids colored by A) time points and B) species (Aha – *A. halleri*, Aly – *A. lyrata*, Ath – *A. thaliana*, CH – F1 *A. thaliana* x *A. halleri*, CM – F1 *A. thaliana* x *A. lyrata*). The PCA separates the samples by species (PC1 59% variance) and by time-points (PC2 18% variance).

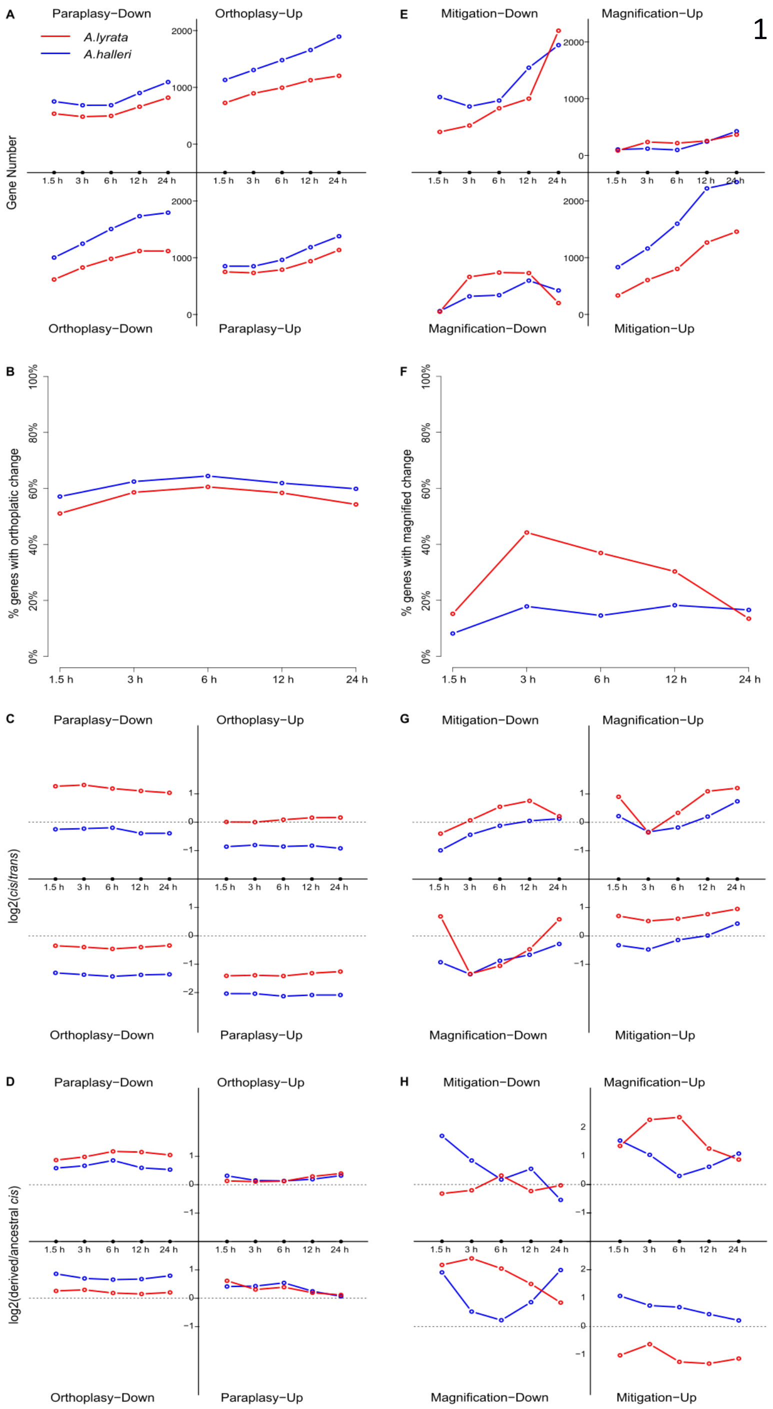


Fig S3. **Extension of Fig. 2 and 3 to all time points of the experiment.** Basal changes (A – D) and plastic responses to the stress (E – H) in *A. lyrata or A. halleri*, compared *to A. thaliana*. (A) The number of genes with a basal change in the direction (ortho-) or opposite (para-) to the direction of the stress reaction in *A. thaliana* increases over the time course. (B) The proportion of genes with an orthoplastic change is not dependent on the time point at which plasticity is determined. (C) The proportion of basal changes explained by *cis*-acting differences is not dependent on the time point at which plasticity is determined. (D) The proportion of basal changes explained by a derived *cis* is not dependent on the time point at which plasticity is observed.

E) The number of genes with a mitigated response increase during the time course. The number of genes with a magnified response do not increase to the same extent. (F) In *A. halleri*, the proportion of genes with a magnified response is not dependent on the time point, whereas in *A. halleri* the proportion of magnified genes is increased between 6 and 12 hours. (G ) The proportion of responses to the stress explained by plastic *cis*-acting differences is dependent on the time point in *A. lyrata*. H) The proportion of responses to the stress explained by derived *cis*-acting changes is dependent on the time point.

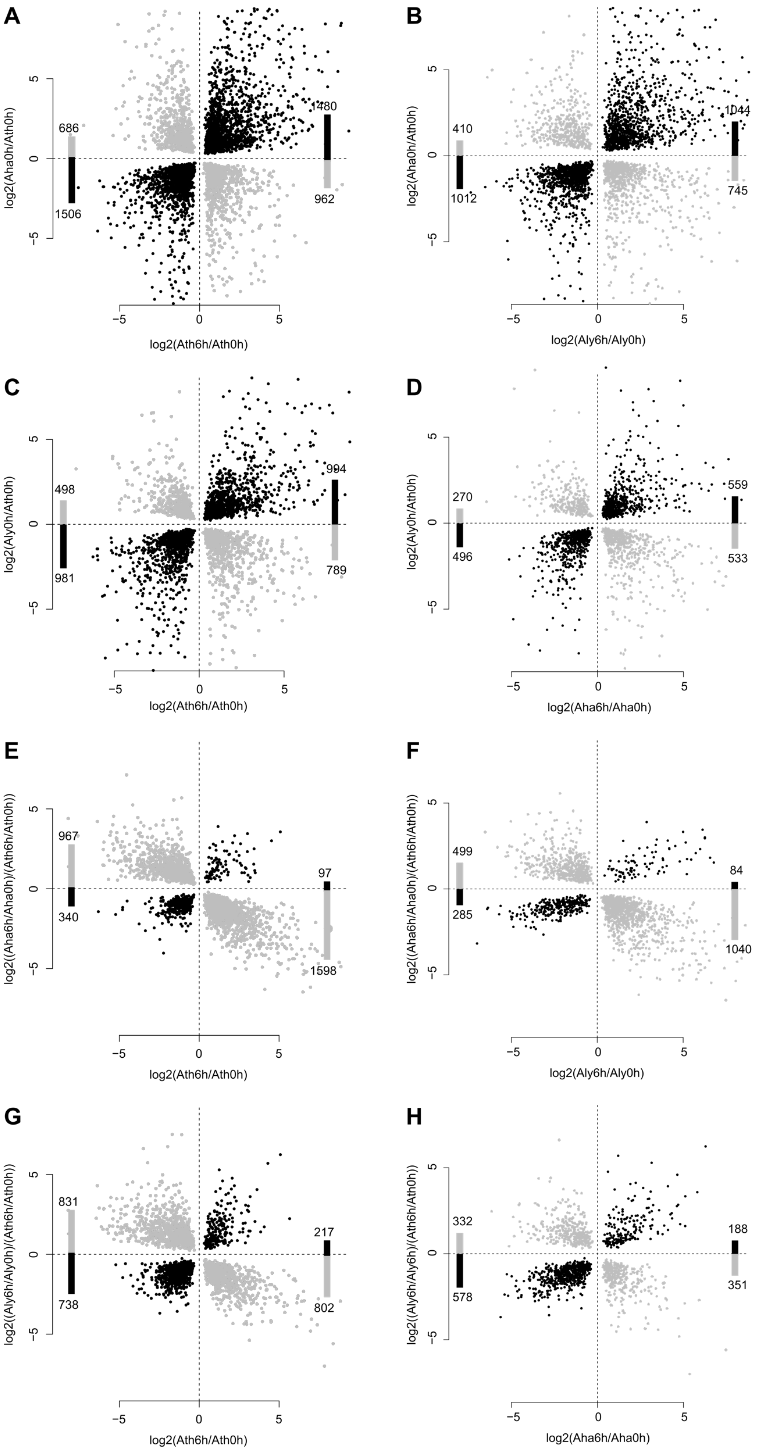


**Figure S4. Orthoplastic evolution of basal (A – D) gene expression in *A. lyrata* or *A. halleri* is robust to the species used for comparison. A mitigated response to stress in *A. halleri* is robust to the proxy used for plastic changes (E – F) . This is not the case in *A. lyrata* (G-H).**

A-B: Basal changes in *A. halleri* vs *A. thaliana* using *A. thaliana* (A) and *A. lyrata* (B) as a comparison.

C-D: Basal changes in *A. lyrata* vs *A. thaliana* using *A. thaliana* (C) and *A. halleri* (D) as a comparison.

E-F: Plastic changes in *A. halleri* using *A. thaliana* (E) and *A. lyrata* (F) as a comparison.

G-H: Plastic changes in *A. lyrata* using *A. thaliana* (E) and *A. halleri* (F) as a comparison.

A-D: orthoplastic: black, paraplastic: grey

E-H: magnification: black, mitigation: grey

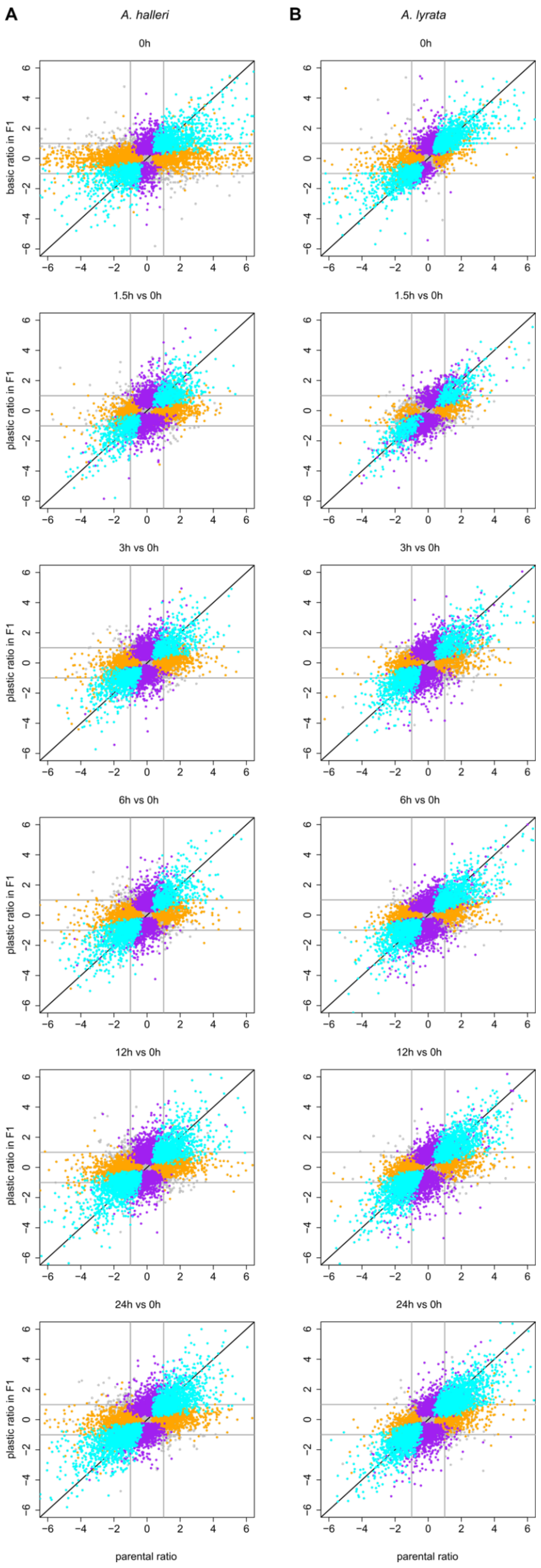


Fig. S5. ***Cis* regulatory changes explain a large part of the interspecific differences between the species. (**A) *A. halleri*. (B) *A. lyrata*. Cyan: *cis* change explains most of the parental difference; Orange parental difference are not associated with a bias in allele specific expression and thus assumed to be controlled in trans- (no significant *cis*-acting change); purple: Significant cis-acting changes are detected but the difference between parents is not significant. The allele-specific expression ratio of purple genes is however strongly correlated with the parental expression ratio, indicating that this class of genes reflect lower power for detecting parental differences (shown in Table S8).

At 0h, the genetic basis of regulatory differences in basal expression is shown by plotting log2ratio of *A. halleri*/*A. thaliana* or *A. lyrata*/*A. thaliana* of alleles in the hybrid (y-axis) against the ratio of parental expression (x-axis). From 1.5h to 24h, the genetic basis of regulatory changes is determined by comparing ratio of log2 ratios at xh vs. 0h for *A. halleri*/*A. thaliana* or *A. lyrata*/*A. thaliana* alleles in hybrid (y-axis) with parents (x-axis) at each time point.

Pearson correlations for *cis* (cyan), non-*cis* (trans; orange) and purple (compensatory; *cis*-change but no significant parental difference) are shown in Table S8.

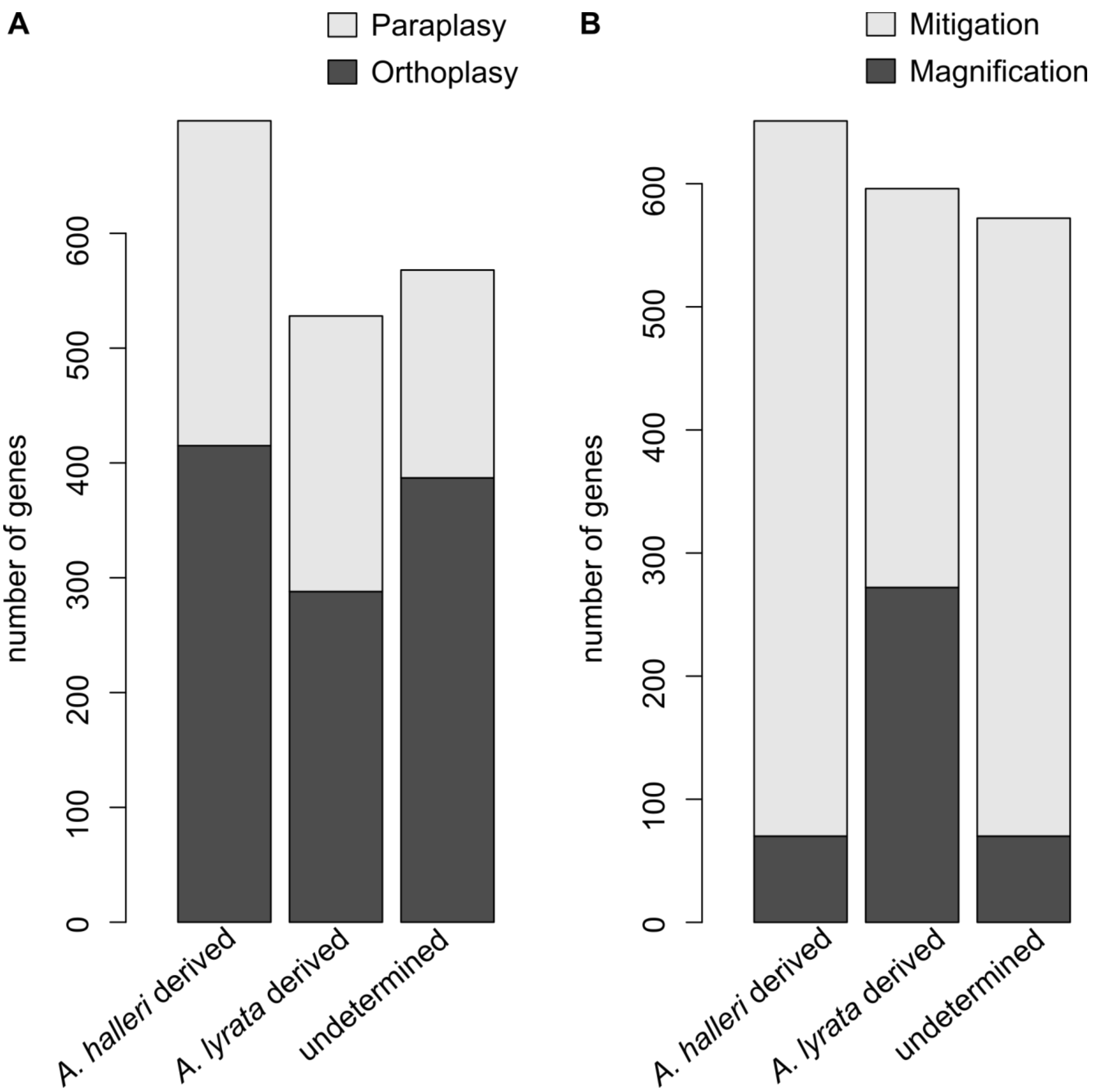


Figure S6. **The total number of genes with derived and undetermined *cis*-regulatory changes are similar.** The number of genes that have (A) a basal change and either an undetermined or derived *cis*-regulatory change or a (B) plastic response and either an undetermined or derived *cis*-regulatory change. Genes are partitioned by the direction of the cis-acting effect detected in basal expression (A) or in the response slope (B). The proportions from Fig4C are based on these numbers. Because the direction of effect of cis-regulatory changes of undetermined origin cannot be ascertained, the number of genes with these changes are not partitioned functionally.

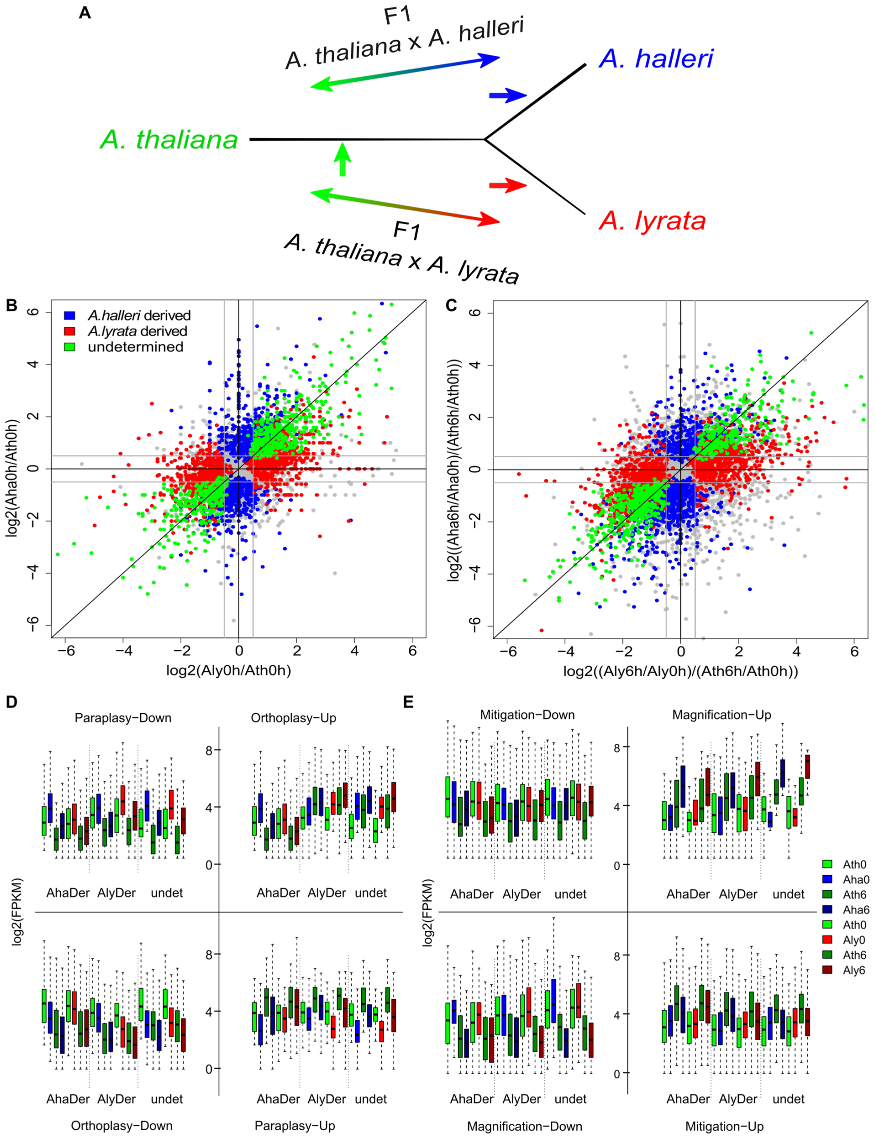


Fig. S7. **The phylogenetic origin of cis-acting changes was determined based on the comparison of allele-specific expression ratios in the two F1 hybrids *A. thaliana-A. lyrata* and *A. thaliana-A. halleri*.** (A) Phylogenetic relationship between the three species allowed distinguishing cis-acting changes of undetermined origin (green arrow), which are shared in the two F1 hybrids, from the derived *cis*-acting changes, which are specific (blue and red arrow). Also shown in Fig. 4B. (B) basal and (C) plastic cis-acting changes. Log2 allelic ratios in AthxAha hybrids plotted against log2 allelic ratios in AthxAly hybrids in basal cis mutations (B), genes with an undetermined *cis*-regulatory change are on the diagonal (green), genes with a derived *cis*-regulatory change in *Arabidopsis lyrata* are on the horizontal central lines (red) and genes with a derived *cis*-regulatory change in *A. halleri* are on the vertical line (blue). Log2 of allelic ratio changes between 6h and 0h in AthxAha hybrids plotted against log2 allelic ratio changes in AthxAly hybrids in plastic cis mutations (C). Box and whiskers depict the 75th and 95$^{th}$ interquantile ranges, respectively, dots show outliers.

**The inferred phylogenetic inference of derived *cis*-acting changes is independent of read count levels.** D). The total expression level before stress for the orthoplastic and paraplastic genes in the F1 hybrids. Dark colors are for time point 6h, light colors for time point 0 hours. *A. thaliana*: green, *A. halleri*: blue and *A. lyrata*: red. E). The total expression level before and after stress for the magnified and mitigated genes in the F1 hybrids. Dark colors are for time point 6h, light colors for time point 0 hours. *A. thaliana*: green, *A. halleri*: blue and *A. lyrata*: red.

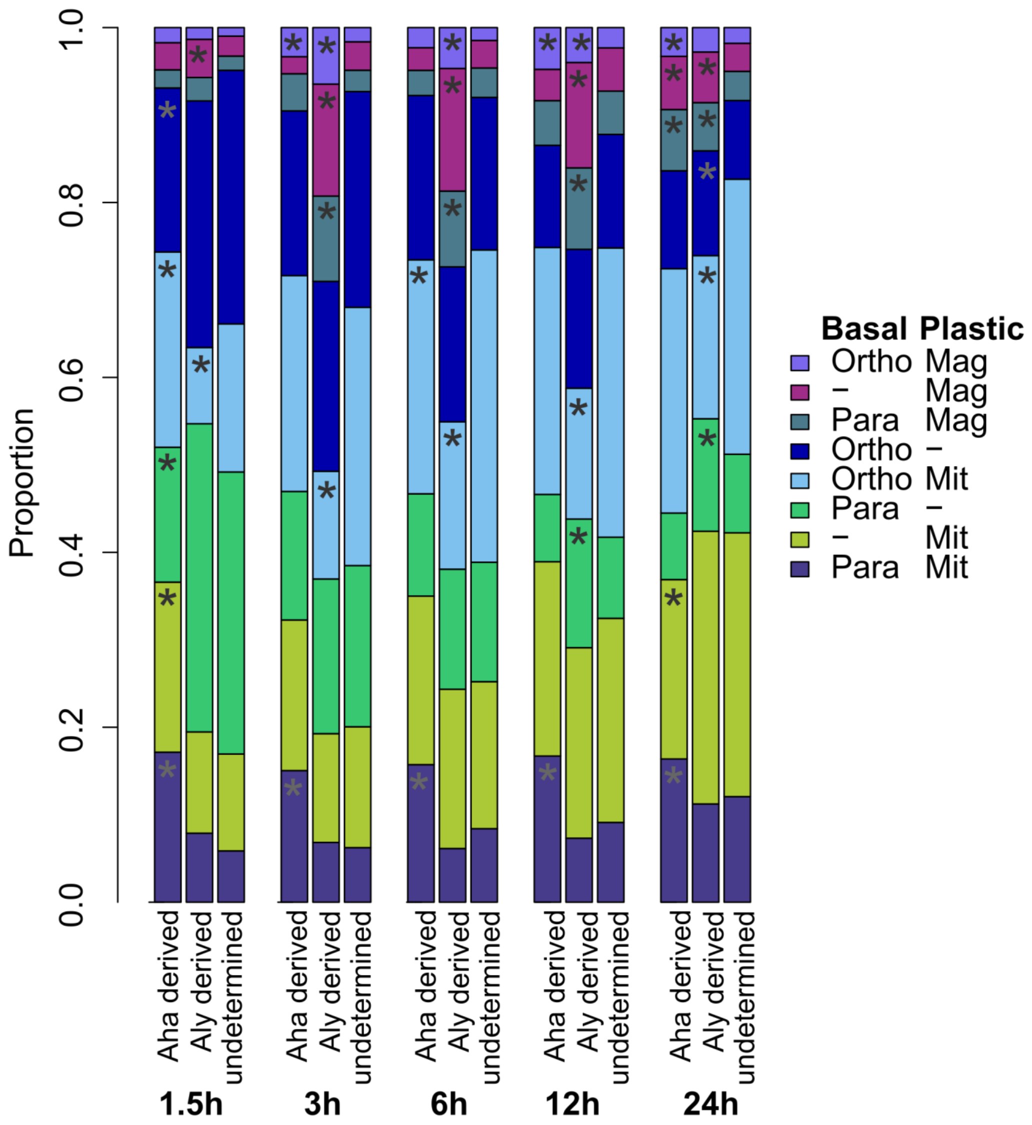


Figure S8. **The pattern reported in Fig. 4C is also apparent in samples collected after 3h or 12h.**

Ortho-Mag: orthoplasy and magnification, Ortho-Mit: orthoplasy and mitigation, Para-mit: paraplasy and mitigation, Para-Mag: paraplasy and magnification, Only-Mag: magnification only, Only-mit: Mitigation only, Only-Ortho: Orthoplasy only, Only-Para: Paraplasy only.

The proportion of derived and undetermined *cis*-regulatory modifications observed after 6h within each class of genes combining basal and/or plastic expression changes is similar after 3h or 12h of desiccation stress, at the time points where the largest numbers of genes are observed that show a magnified response to stress. Aha - *A. halleri,* Aly– *A. lyrata*.

"*": Significantly increased or decreased number of derived compared to undetermined changes in *A. halleri* or *A. lyrata* at each time point.

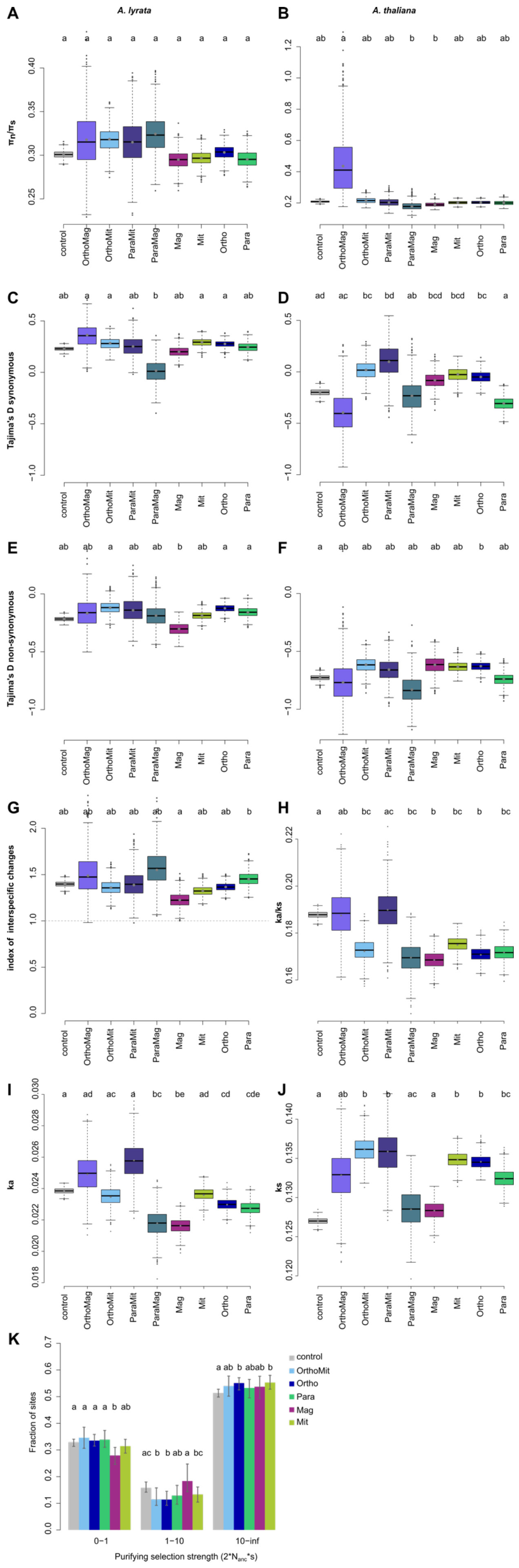


Fig. S9: Summary statistics for the 8 plasticity groups and the control for *A. lyrata* (A,C,E) and *A. thaliana* (B,D,F). Significance between the groups was estimated based on pairwise comparison of 1000 bootstrap replicates, by using the union of the bootstrap values to calculate what proportion of the differences between the groups significantly differed from 0, multiplied by 2, to account for the fact that the test is one-sided. Box and whiskers depict the 75th and 95$^{th}$ interquantile ranges, respectively, dots show outliers. No shared letters mean that the groups are different with a significance cutoff at $p < 0.05$. Pvalues of all pairwise comparisons can be found in Supplementary data 7.

A) and B) pi$_n$/pi$_s$, C and D) Tajima's D for synonymous sites, E and F) Tajima's D for non-synonymous sites. G) index of interspecific change for nonsynonymous sites ((A. lyrata +2) / ((A. thaliana +2)), H) ka/ks between *A. thaliana* and *A. lyrata,* I) ka and J) ks.

The plasticity classes OrthoMag, ParaMit and ParaMag were removed from analysis due to the low number of genes and the high variance between the bootstrap in these categories.

K) binned DFE for the control and 5 of the plasticity classes. 95% confidence intervals around the observed mean are based on 200 bootstrap replicates and depicted by error bars. P-values can be found in Table S9.

No shared letters between plasticity groups indicates a significant difference between the groups.

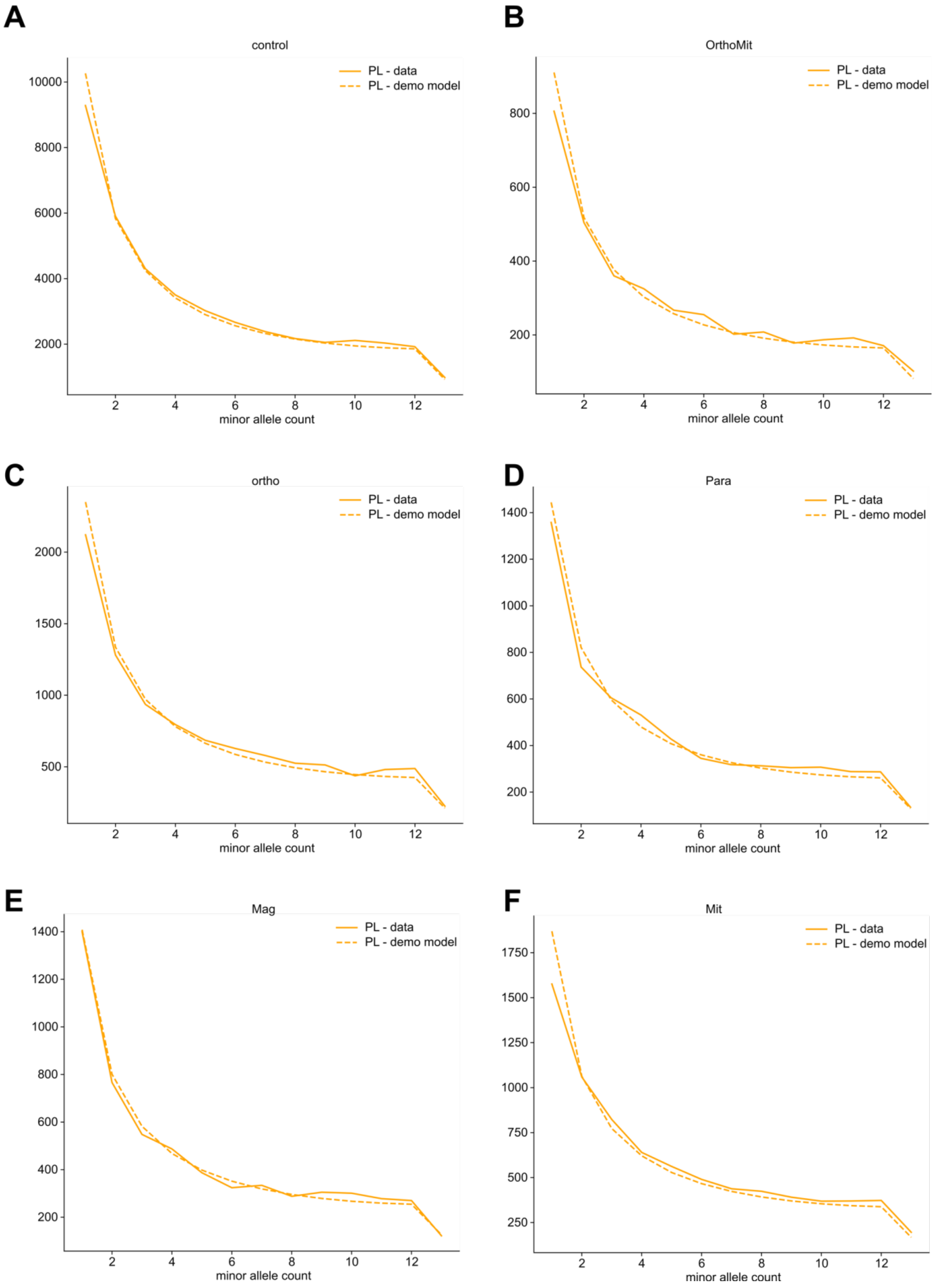


Figure S10. Folded site frequency spectra for synonymous sites based on the data (solid line) and based on the expected number of sites of the demographic model (dashed line) for 5 sufficiently large gene sets, that grouped genes by their mode of plasticity evolution, as well as for the group of control genes, which had the same expression levels at all time points in all three species.

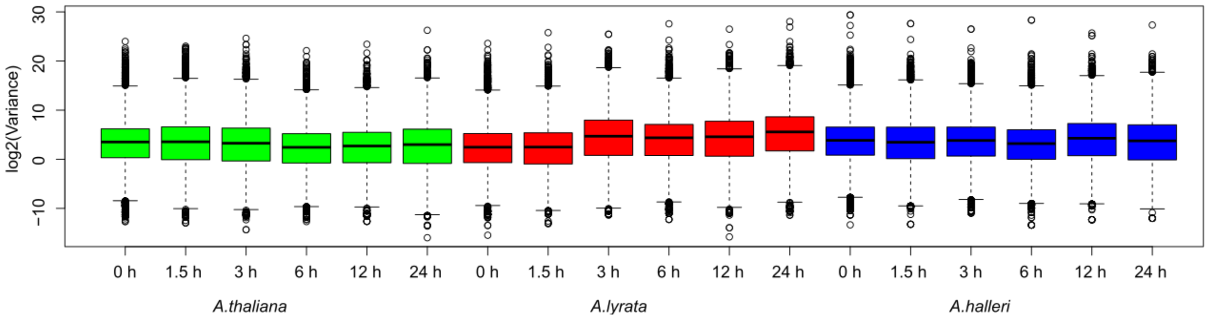


Fig. S11. **The variance of gene expression among biological replicates**. The variance of the expression level in the samples does not increase with the duration of exposure to the stress. The variance of each gene, in each species and at each time point, was calculated based on the four independent biological replicates of the experiment. Box and whiskers depict the 75th and 95$^{th}$ interquantile ranges, respectively; dots show outliers.

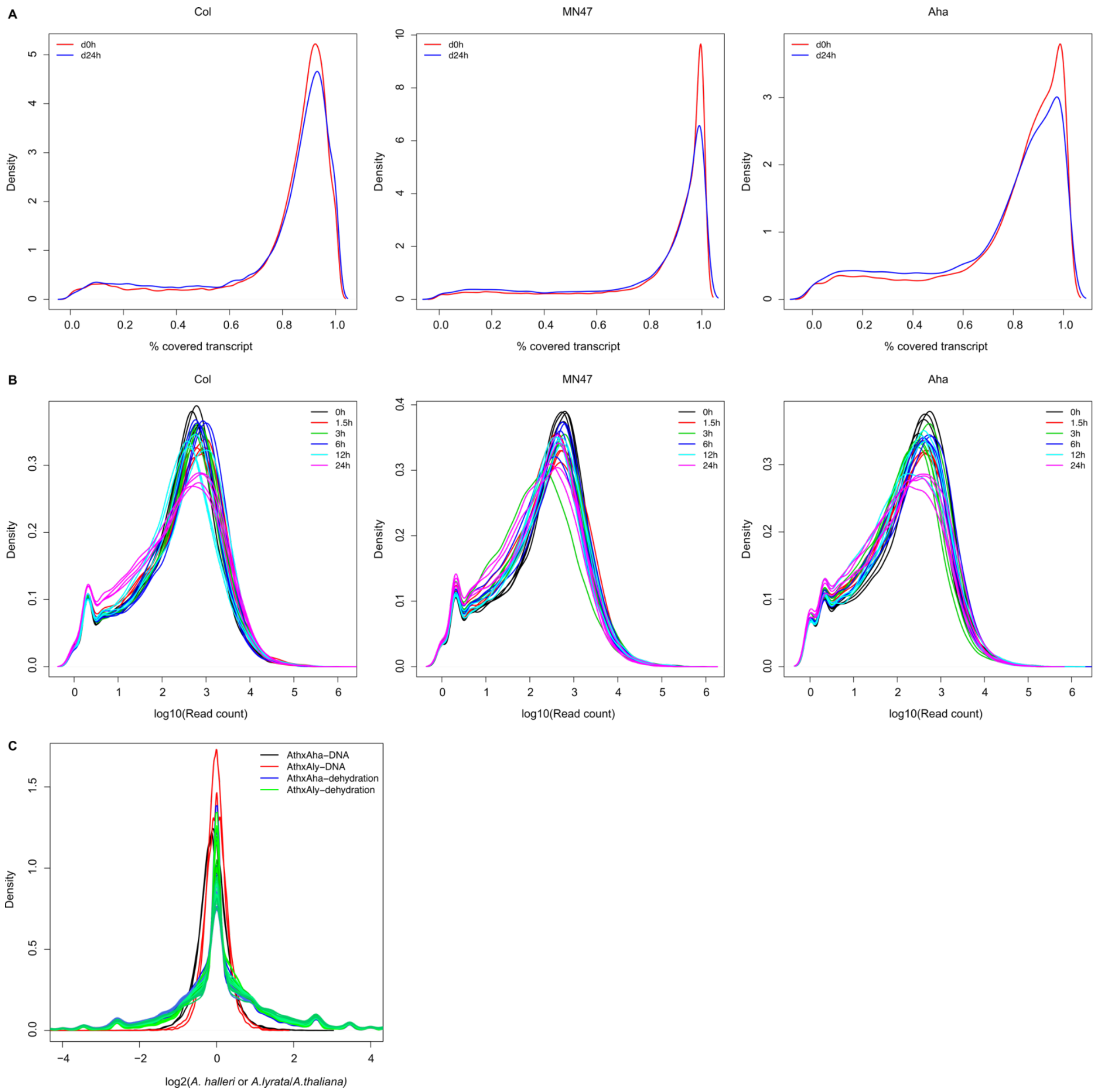


Figure S12. **Data quality is comparable across samples.** (A) The proportion of the covered transcript at time point 0 hours (red) and time point 24 hours (blue) for the three species. (B) The distribution of the log10 read count at all time points (0 hours – 24 hours) for the three species and (C) the distribution of the log2 allele ratio in the F1s after remapping and low divergent region filter for the genomes (AthxAha – black; AthxAly – red) and the transcriptomes (AthxAha – blue; AthxAly – green).

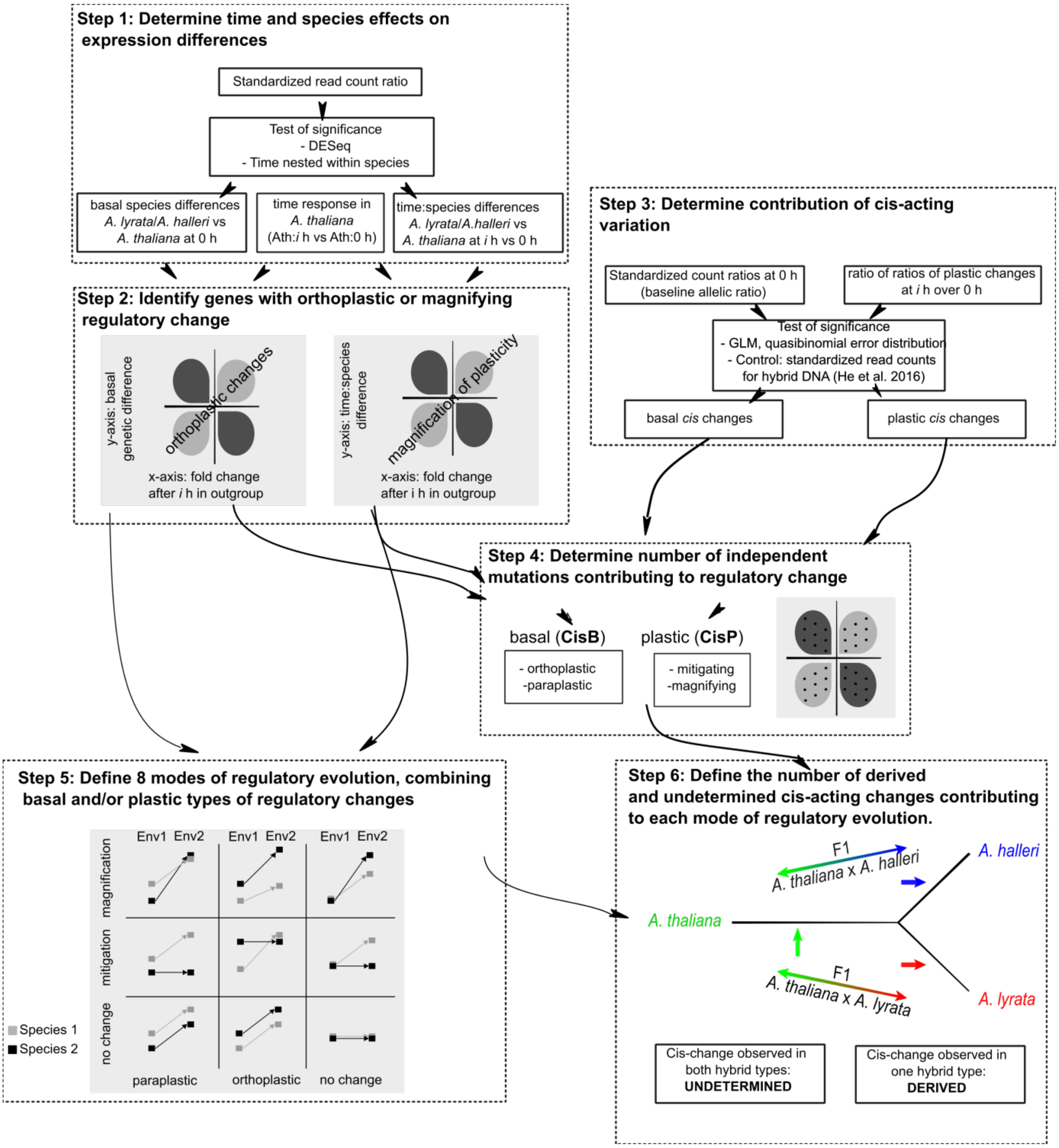


Figure S13. Overview of the analytical pipeline to determine basal and plastic expression changes (step 1, step 2) and infer the number of independent *cis*-changes (Step 3, Step 4). Step 5: integration of analyses describing basal and plastic changes in gene regulation in *A. lyrata* and *A. halleri* to define 8 modes of plasticity evolution as well as a group of genes that do not show any differences. Step 6: inferring the derived state of the *cis*-changes observed.

Table S1: Percentage of differentially expressed genes that overlap with the differentially expressed genes in Bouzid et al. (2019) for A. halleri and A. lyrata. The study included three different time points, 60% soil moisture (before the stress starts), 20% soil moisture (dehydration stress) and recovery (time point after rewatering). The percentage of up and down regulated in this study is in the top row. Significant overlap between the studies was tested using a hypergeometric test.

Table S1

| | | A. halleri 1 vs 0 h 22% up, 19% down | A. halleri 3 vs 0 h 27% up, 24.9% down | A. halleri 6 vs 0 h 26.5% up, 24.7% down | A. halleri 12 vs 0 h 32.6% up, 30.8% down | A. halleri 24 vs 0 h 35% up, 34.8% down |
|---|---|---|---|---|---|---|
| A. halleri 20 % vs 60 % soil moisture | Up (127 ATG genes) | 33% p= 0.001815975 | 37% p = 0.003911163 | 44% p= 6.16E-06 | 57% p= 2.48E-09 | 63.7% p=9.60E-12 |
| | Down (385 ATG genes) | 35% p=1.23E-14 | 42.4% p=6.33E-11 | 54% p=7.49E-37 | 61% p=2.87E-36 | 64% p=4.56E-34 |

| | | A. halleri 1 vs 0 h 22% up, 19% down | A. halleri 3 vs 0 h 27% up, 24.9% down | A. halleri 6 vs 0 h 26.5% up, 24.7% down | A. halleri 12 vs 0 h 32.6% up, 30.8% down | A. halleri 24 vs 0 h 35% up, 34.8% down |
|---|---|---|---|---|---|---|
| A. halleri recovery vs 60 % soil | Up (6 ATG genes) | 0 n.s. | 0 n.s. | 0 n.s. | 0 n.s. | 0 n.s. |
| | Down ( 7 ATG genes) | 28% n.s. | 42.8% p= 0.03514445 | 28% n.s. | 14% n.s. | 14% n.s. |

| | | A. lyrata 1 vs 0 h 21.8% up; 22.5% down | A. lyrata 3 vs 0 h 32% up, 29% down | A. lyrata 6 vs 0 h 32.8% up, 30.9% down | A. lyrata 12 vs 0 h 35% up, 32% down | A. lyrata 24 vs 0 h 37% up, 31% down |
|---|---|---|---|---|---|---|
| A. lyrata 20 % vs 60 % soil moisture | Up (15 ATG genes) | 40% p= 0.0288373 | 46% n.s. | 53% p= 0.02819508 | 53% p= 4.51E-02 | 53% n.s. |
| | Down (37 ATG genes) | 18.9% n.s. | 13.5% n.s. | 13.5% n.s. | 18.9% n.s. | 16.2% n.s. |

| | | A. lyrata 1 vs 0 h 21.8% up; 22.5% down | A. lyrata 3 vs 0 h 32% up, 29% down | A. lyrata 6 vs 0 h 32.8% up, 30.9% down | A. lyrata 12 vs 0 h 35% up, 32% down | A. lyrata 24 vs 0 h 37% up, 31% down |
|---|---|---|---|---|---|---|
| A. lyrata recovery vs 60 % soil moisture | Up (61 ATG genes) | 32.7% p= 0.01640409 | 29.5% n.s. | 34.4% p= 5.94947E-40 | 34.4% n.s. | 36% n.s. |
| | Down (90 ATG genes) | 27.7% n.s. | 44.4% p= 8.54E-04 | 44.4% p= 2.25E-03 | 61% p= 4.49E-09 | 51% p= 2.26E-05 |

Table S2: Proportion of stress-response genes in each species whose basal expression or slope (plastic difference) differs from another species. For each species, we counted the number of genes showing stress response comparing expression at tpx with tp0 (FDR 0.05), and then calculated how many of them showed an basal gene expression difference between species at tp0 (species1-tp0/species2-tp0) or a species differences in the slope of the plastic reaction between tpx and tp0 ((species1/species2-tpx)/ (species1/species2-tp0) ) at FDR 0.05.

Table S2

| | Aha | | | | Aly | | | | Ath | | | |
|---|---|---|---|---|---|---|---|---|---|---|---|---|
| | basal difference | | plastic difference | | basal difference | | plastic difference | | basal difference | | plastic difference | |
| Time point | Aha vs Ath | Aha vs Aly | Aha vs Ath | Aha vs Aly | Aly vs Ath | Aly vs Aha | Aly vs Ath | Aly vs Aha | Ath vs Aha | Ath vs Aly | Ath vs Aha | Ath vs Aly |
| 1.5h | 56.59% | 52.75% | 30.27% | 11.36% | 40.05% | 52.68% | 12.78% | 13.62% | 55.34% | 38.98% | 30.02% | 13.08% |
| 3h | 55.88% | 52.79% | 34.75% | 17.74% | 39.86% | 53.42% | 30.78% | 20.93% | 57.38% | 41.27% | 34.57% | 28.51% |
| 6h | 56.86% | 53.42% | 35.46% | 22.74% | 39.47% | 53.21% | 33.58% | 27.05% | 57.86% | 40.73% | 37.48% | 32.31% |
| 12h | 56.32% | 52.80% | 45.71% | 26.52% | 40.20% | 53.93% | 34.14% | 28.59% | 57.35% | 40.27% | 48.22% | 34.08% |
| 24h | 56.35% | 52.97% | 45.06% | 31.43% | 39.79% | 53.60% | 36.61% | 30.55% | 56.64% | 39.32% | 47.09% | 38.82% |

Table S3: The Pearson's correlation (r) of log ratio of allelic expression difference between hybrid and parents computed separately for genes with a significant cis change  and a significant difference between parents (cis),  a significant difference between parents only (trans only) and a significant cis change in hybrid only (weak cis). At tp 0, the allelic difference is the ratio of expression difference in alleles between species (species1-tp0 / species2-tp0). At tpx, the allelic difference is the ratio of ratio of allelic difference of species between tpx and tp0 (  (species1/spcies2-tpx) / (speices1/species2-tp0) ). *Cis* means both of significant difference in the hybrid and parents with the same direction of changes. *Trans* means that there is no significant cis change in the hybrids. Compensatory means only differences between hybrid alleles not in parents, but they are still correlated to the parents. All correlations are highly significant $p < 2.2e-16$.

Table S3

| | Aha | | | Aly | | |
|---|---|---|---|---|---|---|
| Time point | cis | trans only | weak cis | cis | trans only | weak cis |
| 0h | 0.814 | 0.234 | 0.299 | 0.854 | 0.467 | 0.514 |
| 1.5h | 0.851 | 0.244 | 0.483 | 0.952 | 0.478 | 0.722 |
| 3h | 0.848 | 0.287 | 0.575 | 0.916 | 0.623 | 0.657 |
| 6h | 0.868 | 0.297 | 0.563 | 0.911 | 0.636 | 0.631 |
| 12h | 0.836 | 0.289 | 0.452 | 0.925 | 0.653 | 0.683 |
| 24h | 0.849 | 0.324 | 0.552 | 0.921 | 0.551 | 0.683 |

Table S4. GO enrichment of plastic genes in each mode of plasticity evolution in *A. halleri* and *A. lyrata*. The mode of plasticity is determined by the combination of basal (orthoplasy / Paraplasy) and plastic (Magnification / Mitigation) changes. GO analysis was done using TopGO with the elim method. Enrichment were computed against the reference ensemble of all the expressed orthologous genes of three species. The GO term with less than 2 genes in the column “significant” were removed.

Table S4

| GO.ID | Term | Annotated | Significant | Expected | resultFisher | basal | platic | species |
|---|---|---|---|---|---|---|---|---|
| GO:0001510 | RNA methylation | 116 | 7 | 0.94 | 0.000041 | Orthoplasy | Magnification | A.halleri |
| GO:0006412 | translation | 304 | 8 | 2.46 | 0.0032 | Orthoplasy | Magnification | A.halleri |
| GO:0010440 | stomatal lineage progression | 43 | 3 | 0.35 | 0.005 | Orthoplasy | Magnification | A.halleri |
| GO:0006333 | chromatin assembly or disassembly | 50 | 3 | 0.4 | 0.0076 | Orthoplasy | Magnification | A.halleri |
| GO:0007020 | microtubule nucleation | 52 | 3 | 0.42 | 0.0085 | Orthoplasy | Magnification | A.halleri |
| GO:0007276 | gamete generation | 160 | 5 | 1.29 | 0.0095 | Orthoplasy | Magnification | A.halleri |
| GO:0009220 | pyrimidine ribonucleotide biosynthetic p... | 111 | 9 | 0.87 | 0.0000002 | Orthoplasy | Magnification | A.lyrata |
| GO:0006164 | purine nucleotide biosynthetic process | 215 | 7 | 1.68 | 0.0015 | Orthoplasy | Magnification | A.lyrata |
| GO:0019288 | isopentenyl diphosphate biosynthetic pro... | 188 | 6 | 1.47 | 0.0036 | Orthoplasy | Magnification | A.lyrata |
| GO:0043414 | macromolecule methylation | 394 | 9 | 3.09 | 0.0037 | Orthoplasy | Magnification | A.lyrata |
| GO:0000478 | endonucleolytic cleavage involved in rRN... | 12 | 2 | 0.09 | 0.0038 | Orthoplasy | Magnification | A.lyrata |
| GO:0016570 | histone modification | 294 | 7 | 2.3 | 0.0082 | Orthoplasy | Magnification | A.lyrata |
| GO:0006790 | sulfur compound metabolic process | 534 | 10 | 4.18 | 0.0088 | Orthoplasy | Magnification | A.lyrata |
| GO:0007000 | nucleolus organization | 19 | 2 | 0.15 | 0.0095 | Orthoplasy | Magnification | A.lyrata |
| GO:0006270 | DNA replication initiation | 55 | 17 | 0.78 | 8E-19 | - | Magnification | A.halleri |
| GO:0006275 | regulation of DNA replication | 115 | 19 | 1.64 | 2.3E-15 | - | Magnification | A.halleri |
| GO:0008283 | cell proliferation | 208 | 23 | 2.97 | 1.9E-14 | - | Magnification | A.halleri |
| GO:0051567 | histone H3-K9 methylation | 152 | 19 | 2.17 | 4.4E-13 | - | Magnification | A.halleri |
| GO:0006306 | DNA methylation | 143 | 16 | 2.04 | 1.9E-10 | - | Magnification | A.halleri |
| GO:0009909 | regulation of flower development | 284 | 21 | 4.05 | 6.2E-10 | - | Magnification | A.halleri |
| GO:0034968 | histone lysine methylation | 193 | 27 | 2.75 | 4.1E-08 | - | Magnification | A.halleri |
| GO:0016458 | gene silencing | 298 | 20 | 4.25 | 8.6E-08 | - | Magnification | A.halleri |
| GO:0016572 | histone phosphorylation | 46 | 8 | 0.66 | 2.4E-07 | - | Magnification | A.halleri |
| GO:0010389 | regulation of G2/M transition of mitotic... | 59 | 8 | 0.84 | 0.0000017 | - | Magnification | A.halleri |
| GO:0010075 | regulation of meristem growth | 136 | 11 | 1.94 | 0.0000038 | - | Magnification | A.halleri |
| GO:0000911 | cytokinesis by cell plate formation | 169 | 12 | 2.41 | 0.0000053 | - | Magnification | A.halleri |
| GO:0032508 | DNA duplex unwinding | 10 | 4 | 0.14 | 0.0000078 | - | Magnification | A.halleri |
| GO:0007020 | microtubule nucleation | 52 | 6 | 0.74 | 0.000091 | - | Magnification | A.halleri |
| GO:0051726 | regulation of cell cycle | 240 | 21 | 3.42 | 0.00015 | - | Magnification | A.halleri |
| GO:0006342 | chromatin silencing | 193 | 10 | 2.75 | 0.00044 | - | Magnification | A.halleri |
| GO:0010492 | maintenance of shoot apical meristem ide... | 12 | 3 | 0.17 | 0.00057 | - | Magnification | A.halleri |
| GO:0000724 | double-strand break repair via homologou... | 50 | 5 | 0.71 | 0.0007 | - | Magnification | A.halleri |
| GO:0048451 | petal formation | 58 | 5 | 0.83 | 0.00139 | - | Magnification | A.halleri |
| GO:0048453 | sepal formation | 58 | 5 | 0.83 | 0.00139 | - | Magnification | A.halleri |
| GO:0051225 | spindle assembly | 40 | 4 | 0.57 | 0.00245 | - | Magnification | A.halleri |
| GO:0031047 | gene silencing by RNA | 242 | 10 | 3.45 | 0.00247 | - | Magnification | A.halleri |
| GO:0045786 | negative regulation of cell cycle | 41 | 4 | 0.58 | 0.00268 | - | Magnification | A.halleri |
| GO:0006261 | DNA-dependent DNA replication | 204 | 24 | 2.91 | 0.00319 | - | Magnification | A.halleri |
| GO:0010440 | stomatal lineage progression | 43 | 4 | 0.61 | 0.0032 | - | Magnification | A.halleri |
| GO:0044772 | mitotic cell cycle phase transition | 67 | 10 | 0.96 | 0.00493 | - | Magnification | A.halleri |
| GO:0010051 | xylem and phloem pattern formation | 82 | 5 | 1.17 | 0.00628 | - | Magnification | A.halleri |
| GO:0048768 | root hair cell tip growth | 10 | 2 | 0.14 | 0.00844 | - | Magnification | A.halleri |
| GO:0048829 | root cap development | 10 | 2 | 0.14 | 0.00844 | - | Magnification | A.halleri |
| GO:0008283 | cell proliferation | 208 | 58 | 9.33 | 1E-30 | - | Magnification | A.lyrata |
| GO:0000911 | cytokinesis by cell plate formation | 169 | 46 | 7.58 | 5.8E-24 | - | Magnification | A.lyrata |
| GO:0006275 | regulation of DNA replication | 115 | 38 | 5.16 | 2.5E-23 | - | Magnification | A.lyrata |
| GO:0006270 | DNA replication initiation | 55 | 25 | 2.47 | 1E-19 | - | Magnification | A.lyrata |
| GO:0051567 | histone H3-K9 methylation | 152 | 39 | 6.82 | 1.8E-19 | - | Magnification | A.lyrata |
| GO:0006306 | DNA methylation | 143 | 34 | 6.41 | 4.9E-16 | - | Magnification | A.lyrata |
| GO:0010389 | regulation of G2/M transition of mitotic... | 59 | 22 | 2.65 | 2.7E-15 | - | Magnification | A.lyrata |
| GO:0010103 | stomatal complex morphogenesis | 122 | 29 | 5.47 | 7.4E-14 | - | Magnification | A.lyrata |
| GO:0016572 | histone phosphorylation | 46 | 17 | 2.06 | 4.9E-12 | - | Magnification | A.lyrata |
| GO:0006261 | DNA-dependent DNA replication | 204 | 53 | 9.15 | 3.6E-11 | - | Magnification | A.lyrata |
| GO:0009909 | regulation of flower development | 284 | 39 | 12.74 | 3.7E-10 | - | Magnification | A.lyrata |
| GO:0019288 | isopentenyl diphosphate biosynthetic pro... | 188 | 29 | 8.43 | 4.9E-09 | - | Magnification | A.lyrata |
| GO:0007020 | microtubule nucleation | 52 | 14 | 2.33 | 4.1E-08 | - | Magnification | A.lyrata |
| GO:0009902 | chloroplast relocation | 89 | 18 | 3.99 | 6.3E-08 | - | Magnification | A.lyrata |
| GO:0051225 | spindle assembly | 40 | 12 | 1.79 | 0.0000001 | - | Magnification | A.lyrata |
| GO:0006364 | rRNA processing | 215 | 29 | 9.64 | 0.0000001 | - | Magnification | A.lyrata |
| GO:0010075 | regulation of meristem growth | 136 | 22 | 6.1 | 1.5E-07 | - | Magnification | A.lyrata |
| GO:0009965 | leaf morphogenesis | 176 | 25 | 7.89 | 2.9E-07 | - | Magnification | A.lyrata |
| GO:0000226 | microtubule cytoskeleton organization | 200 | 44 | 8.97 | 9.5E-07 | - | Magnification | A.lyrata |
| GO:0006084 | acetyl-CoA metabolic process | 66 | 14 | 2.96 | 0.000001 | - | Magnification | A.lyrata |
| GO:0007169 | transmembrane receptor protein tyrosine ... | 87 | 16 | 3.9 | 0.0000013 | - | Magnification | A.lyrata |
| GO:0032508 | DNA duplex unwinding | 10 | 6 | 0.45 | 0.0000014 | - | Magnification | A.lyrata |
| GO:0009220 | pyrimidine ribonucleotide biosynthetic p... | 111 | 18 | 4.98 | 0.000002 | - | Magnification | A.lyrata |
| GO:0010027 | thylakoid membrane organization | 162 | 21 | 7.26 | 0.000011 | - | Magnification | A.lyrata |
| GO:0051726 | regulation of cell cycle | 240 | 47 | 10.76 | 0.000011 | - | Magnification | A.lyrata |

| | | | | | | | | |
|---|---|---|---|---|---|---|---|---|
| GO:0008356 | asymmetric cell division | 20 | 7 | 0.9 | 0.000016 | - | Magnification | A.lyrata |
| GO:0042793 | plastid transcription | 66 | 12 | 2.96 | 0.000031 | - | Magnification | A.lyrata |
| GO:0006346 | methylation-dependent chromatin silencin... | 100 | 15 | 4.48 | 0.000037 | - | Magnification | A.lyrata |
| GO:0048451 | petal formation | 58 | 11 | 2.6 | 0.000045 | - | Magnification | A.lyrata |
| GO:0048453 | sepal formation | 58 | 11 | 2.6 | 0.000045 | - | Magnification | A.lyrata |
| GO:0010207 | photosystem II assembly | 139 | 18 | 6.23 | 0.000048 | - | Magnification | A.lyrata |
| GO:0000724 | double-strand break repair via homologou... | 50 | 10 | 2.24 | 0.000061 | - | Magnification | A.lyrata |
| GO:0015995 | chlorophyll biosynthetic process | 107 | 15 | 4.8 | 0.000082 | - | Magnification | A.lyrata |
| GO:0019344 | cysteine biosynthetic process | 158 | 19 | 7.09 | 0.000083 | - | Magnification | A.lyrata |
| GO:0000280 | nuclear division | 186 | 21 | 8.34 | 0.000091 | - | Magnification | A.lyrata |
| GO:0010639 | negative regulation of organelle organiz... | 18 | 6 | 0.81 | 0.000093 | - | Magnification | A.lyrata |
| GO:0010440 | stomatal lineage progression | 43 | 9 | 1.93 | 0.000099 | - | Magnification | A.lyrata |
| GO:0016117 | carotenoid biosynthetic process | 86 | 13 | 3.86 | 0.00011 | - | Magnification | A.lyrata |
| GO:0006468 | protein phosphorylation | 509 | 55 | 22.83 | 0.00013 | - | Magnification | A.lyrata |
| GO:0035304 | regulation of protein dephosphorylation | 114 | 15 | 5.11 | 0.00017 | - | Magnification | A.lyrata |
| GO:0016458 | gene silencing | 298 | 40 | 13.36 | 0.00017 | - | Magnification | A.lyrata |
| GO:0042127 | regulation of cell proliferation | 91 | 13 | 4.08 | 0.0002 | - | Magnification | A.lyrata |
| GO:0006342 | chromatin silencing | 193 | 26 | 8.65 | 0.00022 | - | Magnification | A.lyrata |
| GO:0034968 | histone lysine methylation | 193 | 47 | 8.65 | 0.00026 | - | Magnification | A.lyrata |
| GO:0009637 | response to blue light | 106 | 14 | 4.75 | 0.00027 | - | Magnification | A.lyrata |
| GO:0030154 | cell differentiation | 777 | 56 | 34.84 | 0.00027 | - | Magnification | A.lyrata |
| GO:0031048 | chromatin silencing by small RNA | 96 | 13 | 4.31 | 0.00034 | - | Magnification | A.lyrata |
| GO:0042023 | DNA endoreduplication | 97 | 13 | 4.35 | 0.00038 | - | Magnification | A.lyrata |
| GO:0048449 | floral organ formation | 134 | 22 | 6.01 | 0.00047 | - | Magnification | A.lyrata |
| GO:0009553 | embryo sac development | 184 | 19 | 8.25 | 0.0006 | - | Magnification | A.lyrata |
| GO:0045893 | positive regulation of transcription, DN... | 356 | 30 | 15.96 | 0.00067 | - | Magnification | A.lyrata |
| GO:0016998 | cell wall macromolecule catabolic proces... | 10 | 4 | 0.45 | 0.00068 | - | Magnification | A.lyrata |
| GO:0016126 | sterol biosynthetic process | 117 | 14 | 5.25 | 0.00074 | - | Magnification | A.lyrata |
| GO:0043085 | positive regulation of catA.lyratatic activit... | 104 | 13 | 4.66 | 0.00074 | - | Magnification | A.lyrata |
| GO:0000741 | karyogamy | 35 | 7 | 1.57 | 0.00079 | - | Magnification | A.lyrata |
| GO:0006399 | tRNA metabolic process | 92 | 12 | 4.13 | 0.0008 | - | Magnification | A.lyrata |
| GO:0006655 | phosphatidylglycerol biosynthetic proces... | 58 | 9 | 2.6 | 0.00102 | - | Magnification | A.lyrata |
| GO:0008361 | regulation of cell size | 47 | 8 | 2.11 | 0.00104 | - | Magnification | A.lyrata |
| GO:0048653 | anther development | 72 | 10 | 3.23 | 0.00132 | - | Magnification | A.lyrata |
| GO:0010948 | negative regulation of cell cycle proces... | 12 | 4 | 0.54 | 0.00148 | - | Magnification | A.lyrata |
| GO:0051053 | negative regulation of DNA metabolic pro... | 13 | 4 | 0.58 | 0.00207 | - | Magnification | A.lyrata |
| GO:0006260 | DNA replication | 246 | 64 | 11.03 | 0.00296 | - | Magnification | A.lyrata |
| GO:0043086 | negative regulation of catA.lyratatic activit... | 44 | 7 | 1.97 | 0.00317 | - | Magnification | A.lyrata |
| GO:0006406 | mRNA export from nucleus | 57 | 8 | 2.56 | 0.00369 | - | Magnification | A.lyrata |
| GO:0009658 | chloroplast organization | 202 | 30 | 9.06 | 0.0038 | - | Magnification | A.lyrata |
| GO:0044772 | mitotic cell cycle phase transition | 67 | 25 | 3 | 0.00382 | - | Magnification | A.lyrata |
| GO:0006606 | protein import into nucleus | 84 | 10 | 3.77 | 0.00423 | - | Magnification | A.lyrata |
| GO:0010218 | response to far red light | 84 | 10 | 3.77 | 0.00423 | - | Magnification | A.lyrata |
| GO:0010114 | response to red light | 85 | 10 | 3.81 | 0.0046 | - | Magnification | A.lyrata |
| GO:0019684 | photosynthesis, light reaction | 229 | 29 | 10.27 | 0.00527 | - | Magnification | A.lyrata |
| GO:0007292 | female gamete generation | 117 | 12 | 5.25 | 0.00618 | - | Magnification | A.lyrata |
| GO:1901657 | glycosyl compound metabolic process | 164 | 15 | 7.35 | 0.00689 | - | Magnification | A.lyrata |
| GO:0031400 | negative regulation of protein modificat... | 40 | 6 | 1.79 | 0.00828 | - | Magnification | A.lyrata |
| GO:0009773 | photosynthetic electron transport in pho... | 40 | 6 | 1.79 | 0.00828 | - | Magnification | A.lyrata |
| GO:0071103 | DNA conformation change | 39 | 11 | 1.75 | 0.00829 | - | Magnification | A.lyrata |
| GO:0071840 | cellular component organization or bioge... | 2475 | 223 | 110.99 | 0.00888 | - | Magnification | A.lyrata |
| GO:0010310 | regulation of hydrogen peroxide metaboli... | 123 | 12 | 5.52 | 0.00912 | - | Magnification | A.lyrata |
| GO:0016132 | brassinosteroid biosynthetic process | 80 | 9 | 3.59 | 0.00937 | - | Magnification | A.lyrata |
| GO:0009926 | auxin polar transport | 80 | 9 | 3.59 | 0.00937 | - | Magnification | A.lyrata |
| GO:0006635 | fatty acid beta-oxidation | 137 | 5 | 0.95 | 0.0026 | Paraplasy | Magnification | A.halleri |
| GO:0071472 | cellular response to salt stress | 13 | 2 | 0.09 | 0.0035 | Paraplasy | Magnification | A.halleri |
| GO:0007033 | vacuole organization | 46 | 3 | 0.32 | 0.0039 | Paraplasy | Magnification | A.halleri |
| GO:0019761 | glucosinolate biosynthetic process | 123 | 8 | 1.62 | 0.00022 | Paraplasy | Magnification | A.lyrata |
| GO:0048451 | petal formation | 58 | 5 | 0.76 | 0.00098 | Paraplasy | Magnification | A.lyrata |
| GO:0048453 | sepal formation | 58 | 5 | 0.76 | 0.00098 | Paraplasy | Magnification | A.lyrata |
| GO:0006261 | DNA-dependent DNA replication | 204 | 8 | 2.69 | 0.00564 | Paraplasy | Magnification | A.lyrata |
| GO:0006972 | hyperosmotic response | 193 | 27 | 15.48 | 0.0031 | Orthoplasy | Mitigation | A.halleri |
| GO:0044248 | cellular catabolic process | 1140 | 116 | 91.44 | 0.0037 | Orthoplasy | Mitigation | A.halleri |
| GO:0009813 | flavonoid biosynthetic process | 161 | 23 | 12.91 | 0.0047 | Orthoplasy | Mitigation | A.halleri |
| GO:0010033 | response to organic substance | 1978 | 193 | 158.65 | 0.0052 | Orthoplasy | Mitigation | A.halleri |
| GO:0010264 | myo-inositol hexakisphosphate biosynthet... | 58 | 11 | 4.65 | 0.0058 | Orthoplasy | Mitigation | A.halleri |
| GO:0042128 | nitrate assimilation | 10 | 4 | 0.8 | 0.0058 | Orthoplasy | Mitigation | A.halleri |
| GO:0046482 | Paraplasy-aminobenzoic acid metabolic process | 29 | 7 | 2.33 | 0.0068 | Orthoplasy | Mitigation | A.halleri |
| GO:0016101 | diterpenoid metabolic process | 44 | 9 | 3.53 | 0.0073 | Orthoplasy | Mitigation | A.halleri |
| GO:0098754 | detoxification | 150 | 21 | 12.03 | 0.0085 | Orthoplasy | Mitigation | A.halleri |
| GO:0006820 | anion transport | 393 | 45 | 31.52 | 0.0095 | Orthoplasy | Mitigation | A.halleri |

| | | | | | | | | |
|---|---|---|---|---|---|---|---|---|
| GO:0009749 | response to glucose | 79 | 13 | 6.34 | 0.0097 | Orthoplasy | Mitigation | A.halleri |
| GO:0005983 | starch catabolic process | 15 | 5 | 0.66 | 0.00034 | Orthoplasy | Mitigation | A.lyrata |
| GO:0007623 | circadian rhythm | 137 | 16 | 6.03 | 0.00034 | Orthoplasy | Mitigation | A.lyrata |
| GO:0002213 | defense response to insect | 22 | 5 | 0.97 | 0.00229 | Orthoplasy | Mitigation | A.lyrata |
| GO:0015718 | monocarboxylic acid transport | 16 | 4 | 0.7 | 0.00443 | Orthoplasy | Mitigation | A.lyrata |
| GO:0051187 | cofactor catabolic process | 117 | 12 | 5.15 | 0.00535 | Orthoplasy | Mitigation | A.lyrata |
| GO:0001676 | long-chain fatty acid metabolic process | 17 | 4 | 0.75 | 0.00559 | Orthoplasy | Mitigation | A.lyrata |
| GO:0009408 | response to heat | 215 | 18 | 9.46 | 0.00675 | Orthoplasy | Mitigation | A.lyrata |
| GO:1905039 | carboxylic acid transmembrane transport | 18 | 4 | 0.79 | 0.00694 | Orthoplasy | Mitigation | A.lyrata |
| GO:0042128 | nitrate assimilation | 10 | 3 | 0.44 | 0.00807 | Orthoplasy | Mitigation | A.lyrata |
| GO:0080147 | root hair cell development | 156 | 14 | 6.87 | 0.00893 | Orthoplasy | Mitigation | A.lyrata |
| GO:0006412 | translation | 304 | 122 | 34.96 | 1E-30 | Orthoplasy | - | A.halleri |
| GO:0019288 | isopentenyl diphosphate biosynthetic pro... | 188 | 73 | 21.62 | 1.9E-22 | Orthoplasy | - | A.halleri |
| GO:0006364 | rRNA processing | 215 | 79 | 24.72 | 2E-22 | Orthoplasy | - | A.halleri |
| GO:0010027 | thylakoid membrane organization | 162 | 54 | 18.63 | 1.1E-13 | Orthoplasy | - | A.halleri |
| GO:0001510 | RNA methylation | 116 | 43 | 13.34 | 6.1E-13 | Orthoplasy | - | A.halleri |
| GO:0015995 | chlorophyll biosynthetic process | 107 | 40 | 12.3 | 2.9E-12 | Orthoplasy | - | A.halleri |
| GO:0042254 | ribosome biogenesis | 262 | 101 | 30.13 | 4.3E-10 | Orthoplasy | - | A.halleri |
| GO:0010207 | photosystem II assembly | 139 | 42 | 15.98 | 1.9E-09 | Orthoplasy | - | A.halleri |
| GO:0016226 | iron-sulfur cluster assembly | 83 | 30 | 9.54 | 3.8E-09 | Orthoplasy | - | A.halleri |
| GO:0009658 | chloroplast organization | 202 | 65 | 23.23 | 4E-09 | Orthoplasy | - | A.halleri |
| GO:0045036 | protein targeting to chloroplast | 57 | 24 | 6.55 | 4.1E-09 | Orthoplasy | - | A.halleri |
| GO:0016117 | carotenoid biosynthetic process | 86 | 30 | 9.89 | 1E-08 | Orthoplasy | - | A.halleri |
| GO:0006636 | unsaturated fatty acid biosynthetic proc... | 55 | 23 | 6.32 | 1E-08 | Orthoplasy | - | A.halleri |
| GO:0019344 | cysteine biosynthetic process | 158 | 44 | 18.17 | 1.3E-08 | Orthoplasy | - | A.halleri |
| GO:0009657 | plastid organization | 327 | 109 | 37.6 | 2.6E-08 | Orthoplasy | - | A.halleri |
| GO:0010103 | stomatal complex morphogenesis | 122 | 36 | 14.03 | 5.4E-08 | Orthoplasy | - | A.halleri |
| GO:0009073 | aromatic amino acid family biosynthetic ... | 79 | 27 | 9.08 | 8.9E-08 | Orthoplasy | - | A.halleri |
| GO:0009902 | chloroplast relocation | 89 | 29 | 10.23 | 9.9E-08 | Orthoplasy | - | A.halleri |
| GO:0006399 | tRNA metabolic process | 92 | 36 | 10.58 | 2.5E-07 | Orthoplasy | - | A.halleri |
| GO:0006098 | pentose-phosphate shunt | 155 | 39 | 17.82 | 0.0000015 | Orthoplasy | - | A.halleri |
| GO:0009220 | pyrimidine ribonucleotide biosynthetic p... | 111 | 31 | 12.76 | 0.0000017 | Orthoplasy | - | A.halleri |
| GO:0006655 | phosphatidylglycerol biosynthetic proces... | 58 | 20 | 6.67 | 0.0000035 | Orthoplasy | - | A.halleri |
| GO:0042274 | ribosomal small subunit biogenesis | 11 | 8 | 1.26 | 0.0000036 | Orthoplasy | - | A.halleri |
| GO:0006418 | tRNA aminoacylation for protein translat... | 37 | 15 | 4.25 | 0.000006 | Orthoplasy | - | A.halleri |
| GO:0042793 | plastid transcription | 66 | 21 | 7.59 | 0.0000086 | Orthoplasy | - | A.halleri |
| GO:0016556 | mRNA modification | 89 | 25 | 10.23 | 0.000015 | Orthoplasy | - | A.halleri |
| GO:0009106 | lipoate metabolic process | 27 | 12 | 3.1 | 0.000017 | Orthoplasy | - | A.halleri |
| GO:0009773 | photosynthetic electron transport in pho... | 40 | 15 | 4.6 | 0.000018 | Orthoplasy | - | A.halleri |
| GO:0009695 | jasmonic acid biosynthetic process | 101 | 27 | 11.61 | 0.000018 | Orthoplasy | - | A.halleri |
| GO:0006546 | glycine catabolic process | 37 | 14 | 4.25 | 0.000031 | Orthoplasy | - | A.halleri |
| GO:0019761 | glucosinolate biosynthetic process | 123 | 29 | 14.14 | 0.00011 | Orthoplasy | - | A.halleri |
| GO:0006766 | vitamin metabolic process | 72 | 20 | 8.28 | 0.00012 | Orthoplasy | - | A.halleri |
| GO:0010218 | response to far red light | 84 | 22 | 9.66 | 0.00015 | Orthoplasy | - | A.halleri |
| GO:0009409 | response to cold | 447 | 77 | 51.4 | 0.00016 | Orthoplasy | - | A.halleri |
| GO:0035304 | regulation of protein dephosphorylation | 114 | 27 | 13.11 | 0.00018 | Orthoplasy | - | A.halleri |
| GO:0016122 | xanthophyll metabolic process | 10 | 6 | 1.15 | 0.00032 | Orthoplasy | - | A.halleri |
| GO:0042538 | hyperosmotic salinity response | 124 | 28 | 14.26 | 0.00032 | Orthoplasy | - | A.halleri |
| GO:0006164 | purine nucleotide biosynthetic process | 215 | 41 | 24.72 | 0.00073 | Orthoplasy | - | A.halleri |
| GO:0009637 | response to blue light | 106 | 24 | 12.19 | 0.0008 | Orthoplasy | - | A.halleri |
| GO:0045893 | positive regulation of transcription, DN... | 356 | 61 | 40.94 | 0.00087 | Orthoplasy | - | A.halleri |
| GO:0006417 | regulation of translation | 50 | 14 | 5.75 | 0.00114 | Orthoplasy | - | A.halleri |
| GO:0019216 | regulation of lipid metabolic process | 35 | 11 | 4.02 | 0.00136 | Orthoplasy | - | A.halleri |
| GO:0010155 | regulation of proton transport | 68 | 17 | 7.82 | 0.00143 | Orthoplasy | - | A.halleri |
| GO:0031408 | oxylipin biosynthetic process | 17 | 7 | 1.95 | 0.00179 | Orthoplasy | - | A.halleri |
| GO:0008652 | cellular amino acid biosynthetic process | 351 | 88 | 40.36 | 0.00211 | Orthoplasy | - | A.halleri |
| GO:0048481 | plant ovule development | 120 | 25 | 13.8 | 0.00219 | Orthoplasy | - | A.halleri |
| GO:0009611 | response to wounding | 236 | 42 | 27.14 | 0.00256 | Orthoplasy | - | A.halleri |
| GO:0048316 | seed development | 564 | 90 | 64.86 | 0.0026 | Orthoplasy | - | A.halleri |
| GO:0010675 | regulation of cellular carbohydrate meta... | 18 | 7 | 2.07 | 0.00264 | Orthoplasy | - | A.halleri |
| GO:0010304 | PSII associated light-harvesting complex... | 24 | 8 | 2.76 | 0.00405 | Orthoplasy | - | A.halleri |
| GO:0009414 | response to water deprivation | 293 | 49 | 33.69 | 0.00437 | Orthoplasy | - | A.halleri |
| GO:0016119 | carotene metabolic process | 11 | 5 | 1.26 | 0.00508 | Orthoplasy | - | A.halleri |
| GO:0009965 | leaf morphogenesis | 176 | 32 | 20.24 | 0.00564 | Orthoplasy | - | A.halleri |
| GO:0009867 | jasmonic acid mediated signaling pathway | 211 | 37 | 24.26 | 0.0057 | Orthoplasy | - | A.halleri |
| GO:0043648 | dicarboxylic acid metabolic process | 31 | 9 | 3.56 | 0.00657 | Orthoplasy | - | A.halleri |
| GO:2000377 | regulation of reactive oxygen species me... | 137 | 26 | 15.75 | 0.00676 | Orthoplasy | - | A.halleri |
| GO:0071555 | cell wall organization | 359 | 57 | 41.28 | 0.00695 | Orthoplasy | - | A.halleri |
| GO:0010114 | response to red light | 85 | 18 | 9.77 | 0.00721 | Orthoplasy | - | A.halleri |
| GO:0009863 | salicylic acid mediated signaling pathwa... | 250 | 42 | 28.75 | 0.00731 | Orthoplasy | - | A.halleri |
| GO:0009408 | response to heat | 215 | 37 | 24.72 | 0.00774 | Orthoplasy | - | A.halleri |

| | | | | | | | | |
|---|---|---|---|---|---|---|---|---|
| GO:2000034 | regulation of seed maturation | 12 | 5 | 1.38 | 0.00789 | Orthoplasy | - | A.halleri |
| GO:0006984 | ER-nucleus signaling pathway | 12 | 5 | 1.38 | 0.00789 | Orthoplasy | - | A.halleri |
| GO:0019684 | photosynthesis, light reaction | 229 | 61 | 26.33 | 0.00844 | Orthoplasy | - | A.halleri |
| GO:0044238 | primary metabolic process | 5660 | 812 | 650.87 | 0.00875 | Orthoplasy | - | A.halleri |
| GO:0006413 | translational initiation | 38 | 10 | 4.37 | 0.00902 | Orthoplasy | - | A.halleri |
| GO:0006414 | translational elongation | 17 | 6 | 1.95 | 0.00922 | Orthoplasy | - | A.halleri |
| GO:0009788 | negative regulation of abscisic acid-act... | 22 | 7 | 2.53 | 0.00939 | Orthoplasy | - | A.halleri |
| GO:0009735 | response to cytokinin | 197 | 34 | 22.65 | 0.00993 | Orthoplasy | - | A.halleri |
| GO:0001510 | RNA methylation | 116 | 29 | 9.29 | 2.2E-08 | Orthoplasy | - | A.lyrata |
| GO:0006412 | translation | 304 | 47 | 24.36 | 0.0000088 | Orthoplasy | - | A.lyrata |
| GO:0006635 | fatty acid beta-oxidation | 137 | 24 | 10.98 | 0.00021 | Orthoplasy | - | A.lyrata |
| GO:0009407 | toxin catabolic process | 138 | 23 | 11.06 | 0.00059 | Orthoplasy | - | A.lyrata |
| GO:0043161 | proteasome-mediated ubiquitin-dependent ... | 91 | 17 | 7.29 | 0.0008 | Orthoplasy | - | A.lyrata |
| GO:0009112 | nucleobase metabolic process | 18 | 6 | 1.44 | 0.00209 | Orthoplasy | - | A.lyrata |
| GO:0070417 | cellular response to cold | 13 | 5 | 1.04 | 0.00244 | Orthoplasy | - | A.lyrata |
| GO:0048574 | long-day photoperiodism, flowering | 19 | 6 | 1.52 | 0.00285 | Orthoplasy | - | A.lyrata |
| GO:0009651 | response to salt stress | 571 | 63 | 45.75 | 0.00551 | Orthoplasy | - | A.lyrata |
| GO:0046482 | Paraplasy-aminobenzoic acid metabolic process | 29 | 7 | 2.32 | 0.00677 | Orthoplasy | - | A.lyrata |
| GO:0006221 | pyrimidine nucleotide biosynthetic proce... | 112 | 17 | 8.97 | 0.00767 | Orthoplasy | - | A.lyrata |
| GO:0042402 | cellular biogenic amine catabolic proces... | 70 | 12 | 5.61 | 0.00913 | Orthoplasy | - | A.lyrata |
| GO:0048193 | Golgi vesicle transport | 250 | 31 | 20.03 | 0.00977 | Orthoplasy | - | A.lyrata |
| GO:0009560 | embryo sac egg cell differentiation | 111 | 16 | 7.05 | 0.0017 | Paraplasy | - | A.halleri |
| GO:0006312 | mitotic recombination | 45 | 9 | 2.86 | 0.0018 | Paraplasy | - | A.halleri |
| GO:0009620 | response to fungus | 349 | 36 | 22.15 | 0.0027 | Paraplasy | - | A.halleri |
| GO:0005986 | sucrose biosynthetic process | 12 | 4 | 0.76 | 0.0053 | Paraplasy | - | A.halleri |
| GO:0010200 | response to chitin | 292 | 30 | 18.53 | 0.0062 | Paraplasy | - | A.halleri |
| GO:0019941 | modification-dependent protein catabolic... | 233 | 25 | 14.79 | 0.0069 | Paraplasy | - | A.halleri |
| GO:0006914 | autophagy | 56 | 9 | 3.55 | 0.0082 | Paraplasy | - | A.halleri |
| GO:0048193 | Golgi vesicle transport | 250 | 26 | 15.87 | 0.0089 | Paraplasy | - | A.halleri |
| GO:0019375 | galactolipid biosynthetic process | 77 | 11 | 4.89 | 0.009 | Paraplasy | - | A.halleri |
| GO:0048584 | positive regulation of response to stimu... | 77 | 11 | 4.89 | 0.009 | Paraplasy | - | A.halleri |
| GO:0006914 | autophagy | 56 | 15 | 3.27 | 4.7E-07 | Paraplasy | - | A.lyrata |
| GO:0010200 | response to chitin | 292 | 37 | 17.07 | 0.000007 | Paraplasy | - | A.lyrata |
| GO:0010286 | heat acclimation | 60 | 13 | 3.51 | 0.000033 | Paraplasy | - | A.lyrata |
| GO:0009414 | response to water deprivation | 293 | 38 | 17.12 | 0.000044 | Paraplasy | - | A.lyrata |
| GO:0002679 | respiratory burst involved in defense re... | 74 | 14 | 4.32 | 0.000082 | Paraplasy | - | A.lyrata |
| GO:0009737 | response to abscisic acid | 452 | 50 | 26.42 | 0.00055 | Paraplasy | - | A.lyrata |
| GO:0050832 | defense response to fungus | 234 | 27 | 13.68 | 0.00055 | Paraplasy | - | A.lyrata |
| GO:0002237 | response to molecule of bacterial origin | 64 | 11 | 3.74 | 0.00109 | Paraplasy | - | A.lyrata |
| GO:0035556 | intracellular signal transduction | 338 | 34 | 19.75 | 0.00138 | Paraplasy | - | A.lyrata |
| GO:0009693 | ethylene biosynthetic process | 87 | 13 | 5.08 | 0.00154 | Paraplasy | - | A.lyrata |
| GO:0009697 | salicylic acid biosynthetic process | 142 | 18 | 8.3 | 0.00155 | Paraplasy | - | A.lyrata |
| GO:0042538 | hyperosmotic salinity response | 124 | 16 | 7.25 | 0.0023 | Paraplasy | - | A.lyrata |
| GO:0009620 | response to fungus | 349 | 42 | 20.4 | 0.00232 | Paraplasy | - | A.lyrata |
| GO:0000103 | sulfate assimilation | 11 | 4 | 0.64 | 0.00274 | Paraplasy | - | A.lyrata |
| GO:0009738 | abscisic acid-activated signaling pathwa... | 190 | 21 | 11.1 | 0.00366 | Paraplasy | - | A.lyrata |
| GO:0009753 | response to jasmonic acid | 350 | 33 | 20.46 | 0.00449 | Paraplasy | - | A.lyrata |
| GO:0009627 | systemic acquired resistance | 297 | 29 | 17.36 | 0.0046 | Paraplasy | - | A.lyrata |
| GO:0007267 | cell-cell signaling | 56 | 9 | 3.27 | 0.00483 | Paraplasy | - | A.lyrata |
| GO:0043069 | negative regulation of programmed cell d... | 122 | 15 | 7.13 | 0.00494 | Paraplasy | - | A.lyrata |
| GO:0009863 | salicylic acid mediated signaling pathwa... | 250 | 25 | 14.61 | 0.00606 | Paraplasy | - | A.lyrata |
| GO:0009269 | response to desiccation | 22 | 5 | 1.29 | 0.0077 | Paraplasy | - | A.lyrata |
| GO:0035966 | response to topologically incorrect prot... | 284 | 27 | 16.6 | 0.00863 | Paraplasy | - | A.lyrata |
| GO:0006979 | response to oxidative stress | 339 | 31 | 19.81 | 0.00888 | Paraplasy | - | A.lyrata |
| GO:0016197 | endosomal transport | 24 | 5 | 0.86 | 0.0014 | Paraplasy | Mitigation | A.halleri |
| GO:0031669 | cellular response to nutrient levels | 205 | 15 | 7.35 | 0.007 | Paraplasy | Mitigation | A.halleri |
| GO:0006974 | cellular response to DNA damage stimulus | 226 | 16 | 8.1 | 0.0075 | Paraplasy | Mitigation | A.halleri |
| GO:0010260 | animal organ senescence | 20 | 7 | 1.08 | 0.000055 | - | Mitigation | A.halleri |
| GO:0006635 | fatty acid beta-oxidation | 137 | 16 | 7.41 | 0.0029 | - | Mitigation | A.halleri |
| GO:0031348 | negative regulation of defense response | 201 | 20 | 10.87 | 0.0062 | - | Mitigation | A.halleri |
| GO:0009267 | cellular response to starvation | 191 | 19 | 10.33 | 0.0075 | - | Mitigation | A.halleri |
| GO:0019375 | galactolipid biosynthetic process | 77 | 10 | 4.16 | 0.0083 | - | Mitigation | A.halleri |
| GO:0065008 | regulation of biological quality | 1156 | 81 | 62.51 | 0.0085 | - | Mitigation | A.halleri |
| GO:0042631 | cellular response to water deprivation | 44 | 7 | 2.38 | 0.0087 | - | Mitigation | A.halleri |
| GO:0010264 | myo-inositol hexakisphosphate biosynthet... | 58 | 10 | 2.83 | 0.00044 | - | Mitigation | A.lyrata |
| GO:0006995 | cellular response to nitrogen starvation | 17 | 5 | 0.83 | 0.00103 | - | Mitigation | A.lyrata |
| GO:0009108 | coenzyme biosynthetic process | 273 | 24 | 13.32 | 0.00384 | - | Mitigation | A.lyrata |
| GO:0097164 | ammonium ion metabolic process | 43 | 7 | 2.1 | 0.00443 | - | Mitigation | A.lyrata |
| GO:0072521 | purine-containing compound metabolic pro... | 304 | 25 | 14.84 | 0.00749 | - | Mitigation | A.lyrata |
| GO:0052541 | plant-type cell wall cellulose metabolic... | 17 | 4 | 0.83 | 0.00804 | - | Mitigation | A.lyrata |
| GO:0006766 | vitamin metabolic process | 72 | 9 | 3.51 | 0.00808 | - | Mitigation | A.lyrata |

| | | | | | | | | |
|---|---|---|---|---|---|---|---|---|
| GO:0009739 | response to gibberellin | 141 | 14 | 6.88 | 0.00895 | - | Mitigation | A.lyrata |

Table S5. GO enrichment of plastic genes with derived or undetermined cis changes in each mode of plasticity evolution in *A. halleri* and *A. lyrata*. The mode of plasticity evolution is determined by the combination of basal (Orthoplasy / Paraplasy) and plastic (Magnification / Mitigation) changes. GO analysis was done using TopGO with the elim method. Enrichment were computed against the reference ensemble of all the expressed orthologous genes of three species. The GO term with less than 2 genes in the column “significant” were removed.

Table S5

| GO.ID | Term | Annotate | Significa | Expecte | resultFisher | basal | plastic | cis |
|---|---|---|---|---|---|---|---|---|
| GO:0001510 | RNA methylation | 116 | 3 | 0.21 | 0.0011 | Orthoplasy | Magnification | A.halleri derived |
| GO:0016572 | histone phosphorylation | 46 | 2 | 0.08 | 0.0031 | Orthoplasy | Magnification | A.halleri derived |
| GO:0006333 | chromatin assembly or disassembly | 50 | 2 | 0.09 | 0.0036 | Orthoplasy | Magnification | A.halleri derived |
| GO:0006636 | unsaturated fatty acid biosynthetic proc... | 55 | 2 | 0.1 | 0.0044 | Orthoplasy | Magnification | A.halleri derived |
| GO:0008283 | cell proliferation | 208 | 3 | 0.38 | 0.006 | Orthoplasy | Magnification | A.halleri derived |
| GO:0009220 | pyrimidine ribonucleotide biosynthetic p... | 111 | 5 | 0.35 | 2.30E-05 | Orthoplasy | Magnification | A.lyrata derived |
| GO:0008654 | phospholipid biosynthetic process | 306 | 5 | 0.96 | 2.50E-03 | Orthoplasy | Magnification | A.lyrata derived |
| GO:0046148 | pigment biosynthetic process | 219 | 4 | 0.69 | 4.70E-03 | Orthoplasy | Magnification | A.lyrata derived |
| GO:0006779 | porphyrin-containing compound biosynthet... | 131 | 3 | 0.41 | 7.90E-03 | Orthoplasy | Magnification | A.lyrata derived |
| GO:0010383 | cell wall polysaccharide metabolic proce... | 177 | 3 | 0.26 | 0.0021 | - | Magnification | A.halleri derived |
| GO:0009736 | cytokinin-activated signaling pathway | 59 | 2 | 0.09 | 0.0034 | - | Magnification | A.halleri derived |
| GO:0042546 | cell wall biogenesis | 268 | 3 | 0.4 | 0.0068 | - | Magnification | A.halleri derived |
| GO:0010207 | photosystem II assembly | 139 | 10 | 1.43 | 1.60E-06 | - | Magnification | A.lyrata derived |
| GO:0006364 | rRNA processing | 215 | 12 | 2.22 | 2.10E-06 | - | Magnification | A.lyrata derived |
| GO:0016998 | cell wall macromolecule catabolic proces... | 10 | 4 | 0.1 | 2.20E-06 | - | Magnification | A.lyrata derived |
| GO:0010103 | stomatal complex morphogenesis | 122 | 9 | 1.26 | 4.50E-06 | - | Magnification | A.lyrata derived |
| GO:0035304 | regulation of protein dephosphorylation | 114 | 8 | 1.17 | 2.20E-05 | - | Magnification | A.lyrata derived |
| GO:0019288 | isopentenyl diphosphate biosynthetic pro... | 188 | 10 | 1.94 | 2.40E-05 | - | Magnification | A.lyrata derived |
| GO:0009965 | leaf morphogenesis | 176 | 9 | 1.81 | 8.40E-05 | - | Magnification | A.lyrata derived |
| GO:0043085 | positive regulation of catalytic activit... | 104 | 7 | 1.07 | 9.70E-05 | - | Magnification | A.lyrata derived |
| GO:0019344 | cysteine biosynthetic process | 158 | 8 | 1.63 | 2.20E-04 | - | Magnification | A.lyrata derived |
| GO:0045893 | positive regulation of transcription, DN... | 356 | 12 | 3.67 | 3.00E-04 | - | Magnification | A.lyrata derived |
| GO:0040034 | regulation of development, heterochronic | 32 | 2 | 0.06 | 0.0015 | Paraplasy | Magnification | A.halleri derived |
| GO:0048767 | root hair elongation | 143 | 3 | 0.26 | 0.0021 | Paraplasy | Magnification | A.halleri derived |
| GO:0009059 | macromolecule biosynthetic process | 2572 | 11 | 4.66 | 0.0026 | Paraplasy | Magnification | A.halleri derived |
| GO:0034968 | histone lysine methylation | 193 | 5 | 1.08 | 4.50E-03 | Paraplasy | Magnification | A.lyrata derived |
| GO:0007000 | nucleolus organization | 19 | 2 | 0.11 | 5.00E-03 | Paraplasy | Magnification | A.lyrata derived |
| GO:0006790 | sulfur compound metabolic process | 534 | 8 | 2.99 | 9.70E-03 | Paraplasy | Magnification | A.lyrata derived |
| GO:0019761 | glucosinolate biosynthetic process | 123 | 2 | 0.14 | 0.0086 | Paraplasy | Magnification | Undetermined |
| GO:0009072 | aromatic amino acid family metabolic pro... | 167 | 11 | 3.14 | 0.00031 | Orthoplasy | Mitigation | A.halleri derived |
| GO:1901606 | alpha-amino acid catabolic process | 110 | 8 | 2.07 | 0.00109 | Orthoplasy | Mitigation | A.halleri derived |
| GO:0010363 | regulation of plant-type hypersensitive ... | 275 | 13 | 5.17 | 0.0021 | Orthoplasy | Mitigation | A.halleri derived |
| GO:0006612 | protein targeting to membrane | 276 | 13 | 5.19 | 0.00217 | Orthoplasy | Mitigation | A.halleri derived |
| GO:0009651 | response to salt stress | 571 | 21 | 10.73 | 0.00254 | Orthoplasy | Mitigation | A.halleri derived |
| GO:0006972 | hyperosmotic response | 193 | 10 | 3.63 | 0.00353 | Orthoplasy | Mitigation | A.halleri derived |
| GO:0010033 | response to organic substance | 1978 | 53 | 37.18 | 0.00386 | Orthoplasy | Mitigation | A.halleri derived |
| GO:0009963 | positive regulation of flavonoid biosynt... | 80 | 6 | 1.5 | 0.0039 | Orthoplasy | Mitigation | A.halleri derived |
| GO:0046474 | glycerophospholipid biosynthetic process | 137 | 8 | 2.57 | 0.00432 | Orthoplasy | Mitigation | A.halleri derived |
| GO:0046885 | regulation of hormone biosynthetic proce... | 18 | 3 | 0.34 | 0.00434 | Orthoplasy | Mitigation | A.halleri derived |
| GO:0009593 | detection of chemical stimulus | 13 | 3 | 0.15 | 0.00038 | Orthoplasy | Mitigation | A.lyrata derived |
| GO:0009612 | response to mechanical stimulus | 37 | 3 | 0.42 | 0.00844 | Orthoplasy | Mitigation | A.lyrata derived |
| GO:0031425 | chloroplast RNA processing | 13 | 2 | 0.15 | 0.00923 | Orthoplasy | Mitigation | A.lyrata derived |
| GO:0015718 | monocarboxylic acid transport | 16 | 3 | 0.17 | 0.00064 | Orthoplasy | Mitigation | Undetermined |
| GO:0006984 | ER-nucleus signaling pathway | 12 | 3 | 0.15 | 0.00041 | Orthoplasy | - | A.halleri derived |
| GO:0006364 | rRNA processing | 215 | 10 | 2.73 | 0.00041 | Orthoplasy | - | A.halleri derived |
| GO:0019288 | isopentenyl diphosphate biosynthetic pro... | 188 | 9 | 2.39 | 0.00065 | Orthoplasy | - | A.halleri derived |
| GO:0045036 | protein targeting to chloroplast | 57 | 5 | 0.72 | 0.00076 | Orthoplasy | - | A.halleri derived |
| GO:0006790 | sulfur compound metabolic process | 534 | 16 | 6.78 | 0.00123 | Orthoplasy | - | A.halleri derived |

| | | | | | | | | |
|---|---|---|---|---|---|---|---|---|
| GO:1901566 | organonitrogen compound biosynthetic pro... | 1464 | 35 | 18.59 | 0.00161 | Orthoplasy | - | A.halleri derived |
| GO:0015931 | nucleobase-containing compound transport | 108 | 6 | 1.37 | 0.00251 | Orthoplasy | - | A.halleri derived |
| GO:0006006 | glucose metabolic process | 198 | 8 | 2.51 | 0.00375 | Orthoplasy | - | A.halleri derived |
| GO:0009658 | chloroplast organization | 202 | 8 | 2.56 | 0.00423 | Orthoplasy | - | A.halleri derived |
| GO:0015994 | chlorophyll metabolic process | 161 | 7 | 2.04 | 0.00444 | Orthoplasy | - | A.halleri derived |
| GO:0006414 | translational elongation | 17 | 2 | 0.1 | 0.0041 | Orthoplasy | - | Undetermined |
| GO:0006721 | terpenoid metabolic process | 208 | 5 | 1.18 | 0.0065 | Orthoplasy | - | Undetermined |
| GO:0042138 | meiotic DNA double-strand break formatio... | 62 | 4 | 0.5 | 0.0015 | Paraplasy | - | A.halleri derived |
| GO:0006312 | mitotic recombination | 45 | 3 | 0.36 | 0.0055 | Paraplasy | - | A.halleri derived |
| GO:0051567 | histone H3-K9 methylation | 152 | 5 | 1.22 | 0.0074 | Paraplasy | - | A.halleri derived |
| GO:0033044 | regulation of chromosome organization | 101 | 4 | 0.81 | 0.0087 | Paraplasy | - | A.halleri derived |
| GO:0007131 | reciprocal meiotic recombination | 101 | 4 | 0.81 | 0.0087 | Paraplasy | - | A.halleri derived |
| GO:0009560 | embryo sac egg cell differentiation | 111 | 6 | 1.03 | 0.00059 | Paraplasy | - | A.lyrata derived |
| GO:0048573 | photoperiodism, flowering | 129 | 5 | 1.2 | 0.00708 | Paraplasy | - | A.lyrata derived |
| GO:0006606 | protein import into nucleus | 84 | 4 | 0.78 | 0.00778 | Paraplasy | - | A.lyrata derived |
| GO:0043687 | post-translational protein modification | 86 | 4 | 0.8 | 0.00844 | Paraplasy | - | A.lyrata derived |
| GO:0000377 | RNA splicing, via transesterification re... | 88 | 4 | 0.82 | 0.00914 | Paraplasy | - | A.lyrata derived |
| GO:0009831 | plant-type cell wall modification involv... | 14 | 2 | 0.06 | 0.0016 | Paraplasy | - | Undetermined |
| GO:0006508 | proteolysis | 595 | 8 | 2.55 | 0.0036 | Paraplasy | - | Undetermined |
| GO:0010118 | stomatal movement | 73 | 3 | 0.31 | 0.0037 | Paraplasy | - | Undetermined |
| GO:0006085 | acetyl-CoA biosynthetic process | 12 | 2 | 0.13 | 0.0071 | Paraplasy | Mitigation | A.halleri derived |
| GO:0009081 | branched-chain amino acid metabolic proc... | 14 | 2 | 0.15 | 0.0097 | Paraplasy | Mitigation | A.halleri derived |
| GO:0034614 | cellular response to reactive oxygen spe... | 28 | 2 | 0.08 | 0.0026 | Paraplasy | Mitigation | Undetermined |
| GO:0006259 | DNA metabolic process | 594 | 6 | 1.62 | 0.0048 | Paraplasy | Mitigation | Undetermined |
| GO:0060249 | anatomical structure homeostasis | 46 | 2 | 0.13 | 0.0069 | Paraplasy | Mitigation | Undetermined |
| GO:0006096 | glycolytic process | 157 | 3 | 0.43 | 0.0087 | Paraplasy | Mitigation | Undetermined |
| GO:0001101 | response to acid chemical | 1084 | 26 | 15.28 | 0.005 | - | Mitigation | A.halleri derived |
| GO:0014070 | response to organic cyclic compound | 516 | 15 | 7.27 | 0.0063 | - | Mitigation | A.halleri derived |
| GO:0031348 | negative regulation of defense response | 201 | 8 | 2.83 | 0.0076 | - | Mitigation | A.halleri derived |
| GO:0009617 | response to bacterium | 382 | 12 | 5.38 | 0.0079 | - | Mitigation | A.halleri derived |
| GO:0007154 | cell communication | 1356 | 30 | 19.11 | 0.0081 | - | Mitigation | A.halleri derived |
| GO:0009627 | systemic acquired resistance | 297 | 10 | 4.19 | 0.0094 | - | Mitigation | A.halleri derived |
| GO:0009725 | response to hormone | 1313 | 29 | 18.51 | 0.0094 | - | Mitigation | A.halleri derived |
| GO:0006766 | vitamin metabolic process | 72 | 6 | 0.91 | 0.0003 | - | Mitigation | A.lyrata derived |
| GO:0009269 | response to desiccation | 22 | 3 | 0.28 | 0.0026 | - | Mitigation | A.lyrata derived |
| GO:0009845 | seed germination | 193 | 8 | 2.45 | 0.0032 | - | Mitigation | A.lyrata derived |
| GO:0009106 | lipoate metabolic process | 27 | 3 | 0.34 | 0.0047 | - | Mitigation | A.lyrata derived |
| GO:0009072 | aromatic amino acid family metabolic pro... | 167 | 7 | 2.12 | 0.0054 | - | Mitigation | A.lyrata derived |
| GO:0031667 | response to nutrient levels | 218 | 8 | 2.77 | 0.0066 | - | Mitigation | A.lyrata derived |
| GO:0009638 | phototropism | 10 | 2 | 0.13 | 0.0067 | - | Mitigation | A.lyrata derived |
| GO:0008610 | lipid biosynthetic process | 720 | 10 | 3.8 | 0.0042 | - | Mitigation | Undetermined |
| GO:0065008 | regulation of biological quality | 1156 | 13 | 6.1 | 0.0066 | - | Mitigation | Undetermined |
| GO:0046488 | phosphatidylinositol metabolic process | 79 | 3 | 0.42 | 0.0083 | - | Mitigation | Undetermined |

Table S6: The proportion of mapped RNAseq reads for the three species and the F1 hybrids with different reference genomes. As the draft genome of *A. halleri* v1.1 in Phytozome 12.1 can not cover the whole transcriptome of *A. halleri*, we used the *A. lyrata* genome as the reference for *A. halleri* and its hybrid. We further generated an *A. halleri* pseudo genome (see method) and used it as a reference for *A. halleri* and its hybrid. The pseudogenome increases the mapping efficiency.

Table S6

| Genotype | Reference | 0h | 1.5h | 3h | 6h | 12h | 24h |
|---|---|---|---|---|---|---|---|
| Ath | A. thaliana | 92% | 92% | 92% | 92% | 91% | 90% |
| Aly | A.lyrata | 90% | 90% | 90% | 89% | 88% | 87% |
| Aha | A.lyrata | 70% | 70% | 70% | 70% | 68% | 67% |
| Aha | Aha pseudo | 90% | 90% | 90% | 90% | 89% | 88% |
| AthXAly | AthxAly gen | 90% | 90% | 90% | 90% | 89% | 88% |
| AthxAha | AthxAly gen | 80% | 80% | 80% | 80% | 79% | 78% |
| AthxAha | AthxAha pse | 89% | 89% | 89% | 89% | 88% | 88% |

Table S7: The gene number and contribution of basal/plastic cis changes for each mode of plasticity evolution. CisB is basal cis change, cisP is plastic cis change, and cisBP is both of basal and plastic change.

Table S7

| Mode of plasticity evolution | A.halleri Genes | cisB | cisP | cisBP | A.lyrata Genes | cisB | cisP | cisBP |
|---|---|---|---|---|---|---|---|---|
| orthoplasy - magnification | 119 | 19 | 35 | 7 | 118 | 37 | 25 | 13 |
| magnification | 209 | - | 74 | - | 629 | - | 222 | - |
| paraplasy - magnification | 109 | 19 | 20 | 29 | 208 | 45 | 45 | 56 |
| orthoplasy - mitigation | 1205 | 99 | 329 | 248 | 679 | 60 | 195 | 230 |
| orthoplasy | 1662 | 393 | - | - | 1178 | 475 | - | - |
| paraplasy | 975 | 252 | - | - | 860 | 315 | - | - |
| paraplasy - mitigation | 564 | 64 | 185 | 49 | 219 | 42 | 58 | 46 |
| mitigation | 796 | - | 406 | - | 735 | - | 442 | - |

Table S8 List of plant populations and their geographical distribution of the dehydration experiment in Figure S1

Table S8

| Species | Population | Accession | Location | Latitude | Longitude |
|---|---|---|---|---|---|
| *A. halleri* | Laut | Laut3 | Germany | 49.23 | 14.04 |
| *A. halleri* | Wall | Wall7 | Germany | 49.73 | 12.91 |
| *A. halleri* | Kowa | Kowa4 | Poland | 50.63 | 16.08 |
| *A. halleri* | Bara | Bara4 | Romania | 45.55 | 22.9 |
| *A. halleri* | Nisu | Nisu6 | Romania | 46.24 | 22.55 |
| *A. halleri* | lita | Lita6 | Slovenia | 46.76 | 15.79 |
| *A. halleri* | Lobn | Lobn5 | Slovenia | 46.69 | 16.16 |
| *A. halleri* | Noss | Noss5 | Italy | 44.28 | 14.96 |
| *A. halleri* | Pais | Pais9 | Italy | 46.05 | 10.24 |
| *A. halleri* | hal4.11 | hal4.11 | Italy | 45.86 | 9.84 |
| *A. lyrata* | SB | SB12 | Germany | 51.31 | 10.55 |
| *A. lyrata* | LF | LF10 | Austria | 47.59 | 15.36 |
| *A. lyrata* | LF | LF2 | Austria | 47.59 | 15.36 |
| *A. lyrata* | NT | NT12 | Germany | 49.31 | 11.32 |
| *A. lyrata* | VH | Vosshütte | Austria | 47.58 | 16.1 |
| *A. lyrata* | | Tannenberg1 | | | |
| *A. lyrata* | VOS | VOS | Austria | 57.58 | 16. Okt |
| *A. lyrata* | Plech | Plech.Rock79l | Germany | 49.37 | 11.3 |
| *A. lyrata* | Plech | Plech61.2a | Germany | 49.37 | 11.3 |
| *A. lyrata* | Plech | Plech91.4a | Germany | 49.37 | 11.3 |
| *A. lyrata* | Plech | Plech92.2a | Germany | 49.37 | 11.3 |
| *A. lyrata* | Plech | PlechC3 | Germany | 49.37 | 11.3 |
| *A. lyrata* | HAS | HAS.120 | Germany | 49.47 | 11.25 |
| *A. lyrata* | HAS | HAS122c | Germany | 49.47 | 11.25 |
| *A. lyrata* | HAS | HAS166b | Germany | 49.47 | 11.25 |
| *A. lyrata* | MN47 | MN47 | US | 44.31 | -85.6 |
| *A. lyrata* | Sky | Sky | Scotland | 57.54 | -6.17 |
| *A. thaliana* | IP-Ara-4 | IP.Ara4 | Spain | 41.7 | -3.68 |
| *A. thaliana* | IP-Cmo-3 | IP.Cmo3 | Spain | 40.05 | -4.65 |
| *A. thaliana* | IP-Hoy-0 | IP.Hoy0 | Spain | 40.4 | -5 |
| *A. thaliana* | IP-lab-7 | IP.Lab7 | Spain | 40.87 | -4.5 |
| *A. thaliana* | Amu-0 | Amu0 | Spain | 42.35 | -3.03 |
| *A. thaliana* | Coy-0 | Coy0 | Spain | 40.44 | -4.27 |
| *A. thaliana* | Gud-3 | Gud3 | Spain | 40.65 | -4.11 |
| *A. thaliana* | Hec-0 | Hec0 | Spain | 42.86 | -0.7 |
| *A. thaliana* | Pdl-0 | Pdl0 | Spain | 43.02 | -5.6 |
| *A. thaliana* | Prd-0 | Prd0 | Spain | 41.14 | -3.68 |
| *A. thaliana* | Som-0 | Som0 | Spain | 41.14 | -3.58 |
| *A. thaliana* | Urd-1 | Urd1 | Spain | 42.27 | -2.98 |
| *A. thaliana* | Val-0 | Val0 | Spain | 42.31 | -3.1 |
| *A. thaliana* | Col.FRI | Col.FRI | NA | NA | NA |

Table S9: Pvalues for pairwise comparisons of the bins of the different plasticity categories and the control genes for the DFE in Figure S9.

Table S9

| bin 0-1 | ortho-mit | ortho | para | mag | mit |
|---|---|---|---|---|---|
| control | 0.43 | 0.61 | 0.61 | **0.01** | 0.41 |
| ortho-mit | NA | 0.76 | 0.82 | **0.01** | 0.25 |
| ortho | | NA | 0.98 | **0.01** | 0.25 |
| para | | | NA | **0.01** | 0.27 |
| mag | | | | NA | 0.09 |

| bin 1-10 | ortho-mit | ortho | para | mag | mit |
|---|---|---|---|---|---|
| control | **0.03** | **0.01** | 0.17 | 0.32 | 0.15 |
| ortho-mit | NA | 0.98 | 0.7 | **0.04** | 0.51 |
| ortho | | NA | 0.55 | **0.01** | 0.38 |
| para | | | NA | 0.09 | 0.81 |
| mag | | | | NA | **0.03** |

| bin 10-inf | ortho-mit | ortho | para | mag | mit |
|---|---|---|---|---|---|
| control | 0.18 | **0.01** | 0.26 | 0.37 | **0.02** |
| ortho-mit | NA | 0.5 | 0.95 | 0.99 | 0.55 |
| ortho | | NA | 0.39 | 0.62 | 0.93 |
| para | | | NA | 0.97 | 0.45 |
| mag | | | | NA | 0.48 |
